# Solving the sex ratio scandal in *Melittobia* wasps

**DOI:** 10.1101/2020.11.16.384768

**Authors:** Jun Abe, Ryosuke Iritani, Koji Tsuchida, Yoshitaka Kamimura, Stuart A. West

## Abstract

The scandalous sex ratio behaviour of *Melittobia* wasps has long posed one of the greatest problems for the field of sex allocation. In contrast to the predictions of theory, and the behaviour of numerous other organisms, laboratory experiments have found that *Melittobia* females do not produce less female-biased offspring sex ratios when more females lay eggs on a patch. We resolve this scandal, by showing that, in nature, females of *M. australica* have sophisticated sex ratio behaviour, where their strategy also depends upon whether they have dispersed from the patch where they emerged. When females have not dispersed, they will be laying eggs with close relatives, which keeps local mate competition high, even with multiple females, and so they are selected to produce consistently female-biased sex ratios. Laboratory experiments mimic these conditions. In contrast, when females disperse, they will be interacting with non-relatives, and so they adjust their sex ratio depending upon the number of females laying eggs. Consequently, females appear to use dispersal status as an indirect cue of relatedness, and whether they should adjust their sex ratio in response to the number of females laying eggs on the patch.

## Introduction

Sex allocation has produced many of the greatest success stories in the fields of behavioural and evolutionary ecology^1–4^. Time and time again, relatively simple theory has explained variation in how individuals allocate resources to male and female reproduction. Hamilton’s local mate competition (LMC) theory predicts that when *n* diploid females lay eggs on a patch, and the offspring mate before the females disperse, that the evolutionary stable proportion of male offspring (sex ratio) is (*n*−1)/2*n*^5^ (Fig. 1). A female-biased sex ratio is favoured to reduce competition between sons (brothers) for mates, and to provide more mates (daughters) for those sons^6–8^. Consistent with this prediction, females of > 40 species produce female-biased sex ratios, and reduce this female bias when multiple females lay eggs on the same patch^9^ (higher *n*; Fig. 1). The fit of data to theory is so good that the sex ratio under LMC has been exploited as a ‘model trait’ to study the factors that can constrain ‘perfect adaptation’ ^4,10–13^.

**Figure 1.**
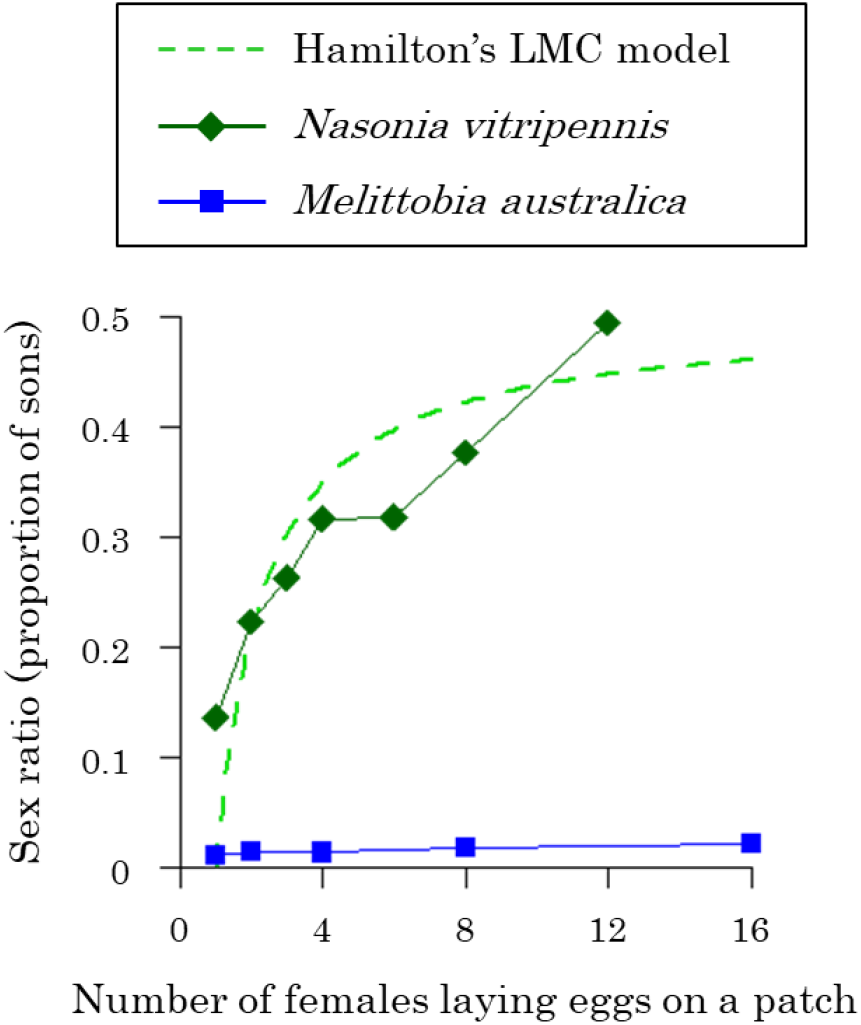
Local mate competition (LMC). The sex ratio (proportion of sons) is plotted versus the number of females laying eggs on a patch. The right green dashed line shows the LMC theory prediction for haplo-diploid species^5,38^. A more female-biased sex ratio is favoured in haplo-diploids^7,31^. Females of many species adjust their offspring sex ratio as predicted by theory, observed in the parasitoid *Nasonia vitripennis*^75^ (green diamonds). In contrast, the females of several *Melittobia* species, such as *M. australica*, continue to produce extremely female-biased sex ratios, irrespective of the number of females laying eggs on a patch^14^ (blue squares).

In stark contrast, the sex ratio behaviour of *Melittobia* wasps has long been seen as one of the greatest problems for the field of sex allocation^3,4,14–21^. The life cycle of *Melittobia* wasps matches the assumptions of Hamilton’s local mate competition theory^5,14,19,21^. Females lay eggs in the larvae or pupae of solitary wasps and bees, and then after emergence, female offspring mate with the short-winged males, who do not disperse. However, laboratory experiments on four *Melittobia* species have found that females lay extremely female-biased sex ratios (1-5% males), and that these extremely female-biased sex ratios change little with increasing number of females laying eggs on a patch^14,17–20,22^ (higher *n*; Fig. 1). A number of hypotheses to explain this lack of sex ratio adjustment have been investigated and rejected, including sex ratio distorters, sex differential mortality, asymmetrical male competition, and reciprocal cooperation^14,16–18,20,22–26^.

We tested whether *Melittobia*’s unusual sex ratio behaviour can be explained by females being related to the other females laying eggs on the same patch. After mating, some females disperse to find new patches, while some may stay at the natal patch to lay eggs on previously unexploited hosts (Fig. 2). If females do not disperse then they can be related to the other females laying eggs on the same host^27–30^. If females laying eggs on a host are related, then this increases the extent to which relatives are competing for mates, and so can favour an even more female-biased sex ratio^28,31–34^. Although most parasitoid species appear unable to directly assess relatedness, dispersal behaviour could provide an indirect cue of whether females are with close relatives^35–37^. Consequently, we predict that when females do not disperse, and so are more likely to be with closer relatives, they should maintain extremely female-biased sex ratios, even when multiple females lay eggs on a patch^28,34^.

**Figure 2.**
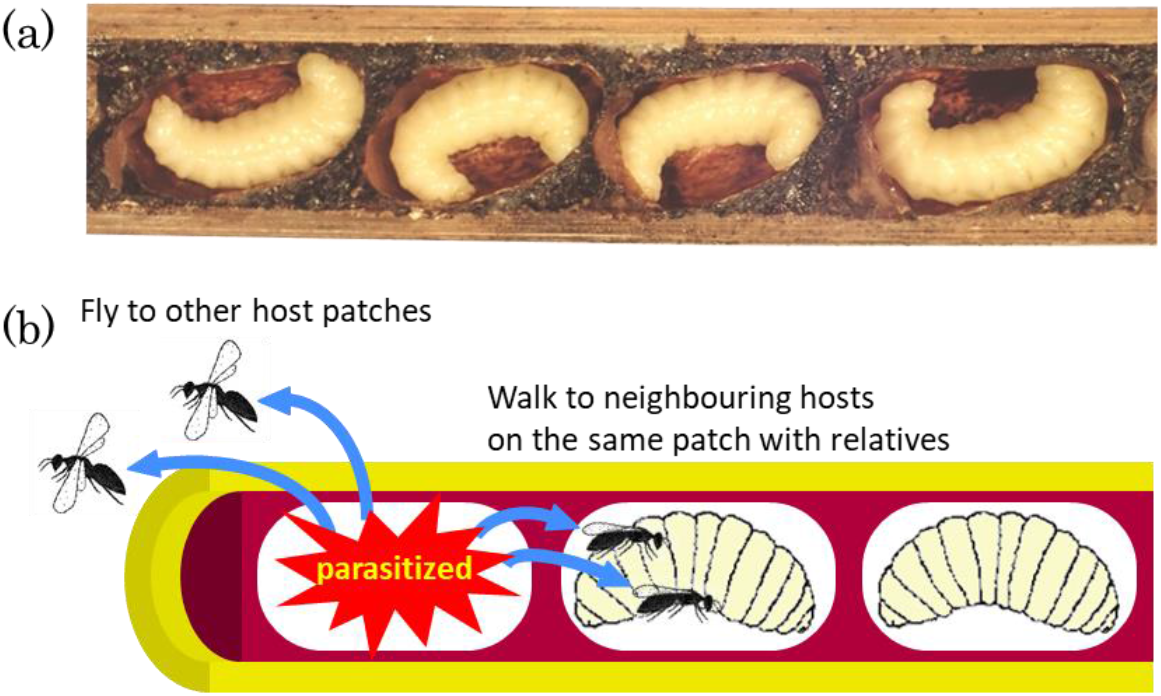
Host nest and dispersal manners of *Melittobia*. (a) Photograph of the prepupae of the leaf-cutter bee *Chalicodoma sculpturalis* nested in a bamboo cane, and (b) diagram showing two ways that *Melittobia* females find new hosts. The mothers of *C. sculpturalis* build nursing nests with pine resin consisting of individual cells, in which their offspring develop. If *Melittobia* wasps parasitize a host in a cell, female offspring that mate with males inside the cell find a different host on the same patch (bamboo cane) or disperse by flying to other patches.

We tested this prediction in a natural population of a *Melittobia* species found in Japan, *M. australica*. We examined how the sex ratio produced by females varies with the number of females laying eggs on a patch, and whether or not they have dispersed before laying eggs. To match our data to the predictions of theory, we developed a mathematical model tailored to the unique population structure of *Melittobia*, where dispersal can be a cue of relatedness. We then conducted a laboratory experiment to test whether *Melittobia* females are able to directly access the relatedness to other females and adjust their sex ratio behaviour accordingly. Our results suggest that females are adjusting their sex ratio in response to both the number of females laying eggs on a patch, and their relatedness to the other females. But, relatedness is assessed indirectly, by whether or not they have dispersed. Consequently, what appeared scandalous, instead reflects a more refined sex ratio strategy.

## Results

### Population structure and relatedness

To obtain the natural broods of *Melittobia*, we placed bamboo traps in the wild (Fig. 2). A total of 4890 host wasps and bees developed in these bamboo traps, with an average of 4.7 ± 2.9 (SD) hosts per bamboo cane (Supplementary Table 2-1). Of these hosts, 0.94% were parasitised by *M. australica*, and we obtained data from 29 *M. australica* broods in which all of the emerging offspring were obtained (Supplementary Table 2-4). We assessed whether the mothers that had laid these broods were from the same host patch (non-dispersers) or had dispersed from a different host patch (dispersers), by examining other parasitised hosts on the same patch (bamboo trap). If there were other parasitised hosts, we also confirmed whether they were dispersers form other patches by genotyping microsatellite DNA markers^27^. Overall, we found that 8 broods were laid by non-dispersering females (5 – 36 mothers), 19 broods were laid by disperseing females (1 – 5 mothers), and 2 broods were laid by mixture of non-dispersers and dispersers (6 mothers).

In nature, we found that 55.2% (16/29) of broods were produced by more than one female. The number of females producing a brood varied from 1 – 36, with an arithmetic mean of 6.7 and a harmonic mean of 1.7 (Supplementary Table 2-2d). Broods where multiple females lay eggs are therefore relatively common. Consequently, the lack of sex ratio adjustment, when multiple females lay eggs on patch, cannot be explained by multiple female broods not occurring or being extremely rare in nature^10^. In the two mixed broods, produced by non-dispersers and dispersers, single dispersers produced all-male clutches, and other females were non-dispersers producing clutches containing both sexes. We carried out analyses below discarding the two all-male clutches, because we were interested in the sex allocation behaviour of mothers producing both sexes. However, we found the same qualitative results irrespective of whether we removed the two broods laid by a mixture of dispersers and non-dispersers (Supplementary Information 1).

Our analysis of relatedness estimated by16 polymorphic microsatellite loci suggests that dispersing females laying eggs on the same patch are unrelated, but that non-dispersing females laying eggs on the same patch are highly related (Supplementary Information 1). As the number of females laying eggs increased, the relatedness between females developing in a brood decreased when the brood was produced by dispersing females, but not when the brood was produced by non-dispersing females (Fig. 3a; dispersers: χ^2^_1_ = 10.15, *P* = 0.001; non-dispersers: χ^2^_1_ = 0.93, *P* = 0.33; interaction between dispersal status and number of females laying eggs: χ^2^ = 12.34, *P* < 0.01). The pattern of relatedness for dispersers closely resembled that expected if unrelated females had produced the broods, while the pattern for non-dispersers was clearly different from this expectation (Fig. 3a).

**Figure 3.**
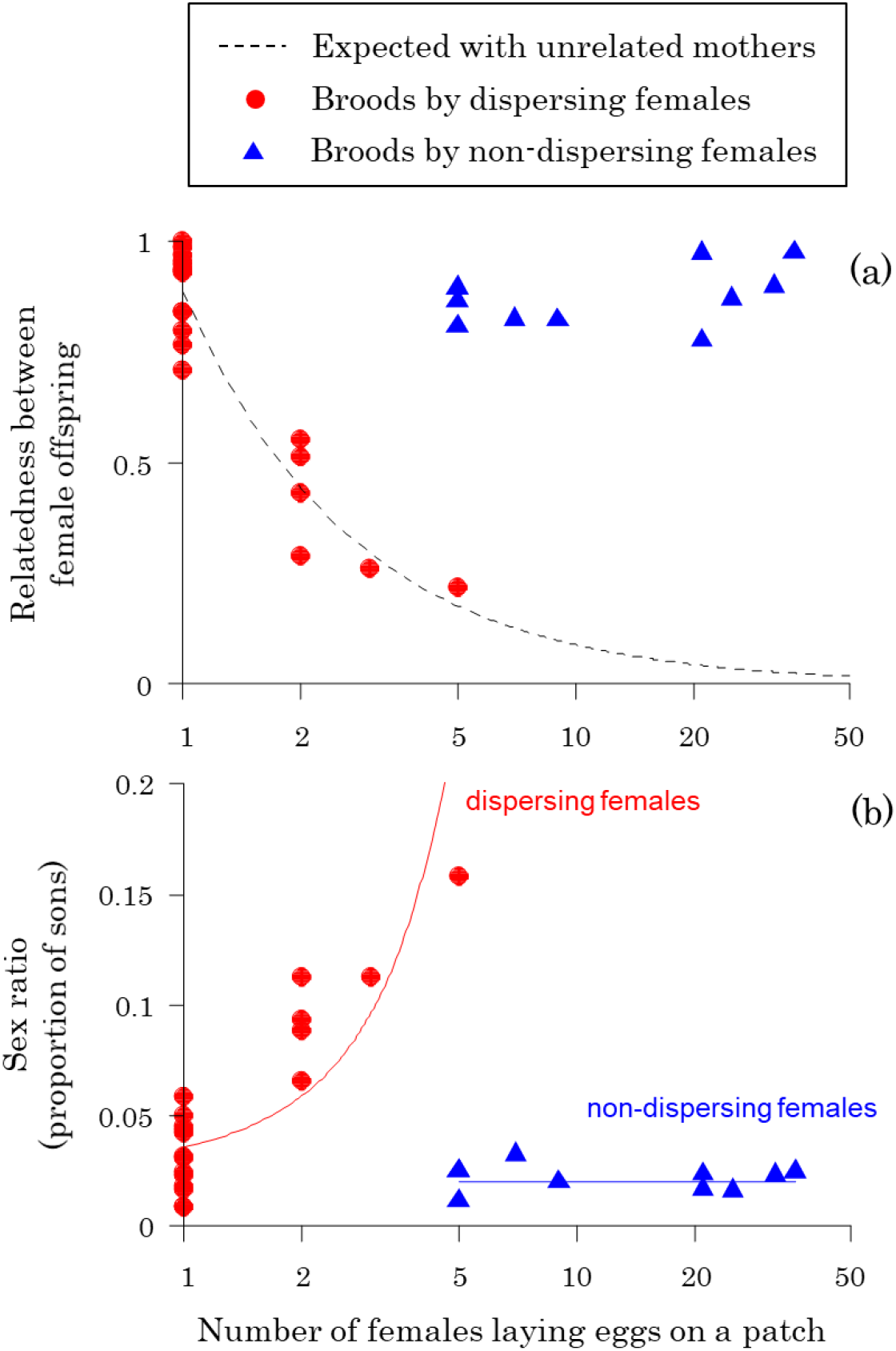
Sex ratios and relatedness in nature. The relationship between the number of females laying eggs on a patch and (a) the average relatedness between female offspring on a patch; (b) the offspring sex ratio (proportion males). The dashed black line in (a) shows the expected relatedness assuming that the mothers of the female offspring are not genetically related, (1 + 3*f*)/(2n(1 + *f*)), where *n* is the number of mothers, and *f* is the inbreeding coefficient estimated as *f* = 0.631 from the microsatellite data. The solid red and blue lines in (b) represent the fitted lines of the generalized linear mixed models assuming binomial distribution for dispersing and non-dispersing females, respectively. The x-axes are logarithmic. When females disperse (red circles), an increasing number of females laying eggs on a patch leads to a decrease in relatedness between female offspring (a) and an increase in the offspring sex ratio (b). In contrast, when females do not disperse, neither the relatedness between female offspring (a) nor the offspring sex ratio (b) significantly varies with the number of females laying eggs on a patch.

The relatedness between the female offspring of non-dispersers remained high regardless of the number of females laying eggs, suggesting that when multiple non-dispersing females laid eggs on the same patch, they were inbred sisters. This was confirmed by our parentage analysis, which suggests that when multiple non-dispersers contributed to a brood, they were offspring of a single female or very closely related females.

### Sex ratios

There was a clear difference in the sex ratios (proportion of sons) produced by dispersing females compared with non-dispersing females. As the number of females laying eggs on a patch increased, dispersers produced less female-biased sex ratios, but non-dispersers did not (Fig. 3b; dispersers: χ^2^_1_ = 14.62, *P* < 0.001; non-dispersers: χ^2^_1_ = 0.56, *P* = 0.46; interaction between dispersal status and number of females laying eggs: χ^2^ = 18.69, *P* < 0.001). Consequently, while non-dispersers always produced approximately 2% males, dispersers produced from 3 to 16% males, as the number of females laying eggs increased.

The pattern in non-dispersers is consistent with laboratory experiments. In laboratory experiments, females were not given a chance to disperse, and as the number of females laying eggs was increased from 1 – 16, there was only a very small increase in offspring sex ratio, from 1 – 2% males^14^ (Fig. 1). In contrast, the increasing sex ratios observed in broods produced by dispersers is consistent with the pattern predicted by Hamilton’s original LMC model^5,38^. The fit to Hamilton’s theory is qualitative, not quantitative, as the observed sex ratios were more female biased.

As also predicted by LMC theory, we found that when individual dispersing females produced more of the offspring on a patch, they produced more female-biased offspring sex ratios^39–41^. The offspring sex ratios produced by a dispersing female was significantly negatively correlated with the proportion of the brood that she produced (Fig. 4; χ^2^_1_ = 9.99, *P* = 0.002). This pattern is predicted by theory, because when a female lays a greater proportion of the offspring on a patch, her sons are more likely to be competing for mates with their brothers, and more likely to be mating with their sisters. Consequently, when females produce a greater proportion of offspring on the patch, their offspring will encounter greater LMC, and so more female-biased sex ratios are favoured^39–41^. Analogous sex ratio adjustment in response to fecundity has been observed in a range of species, including wasps, aphids, and cestodes^39,42–45^.

**Figure 4.**
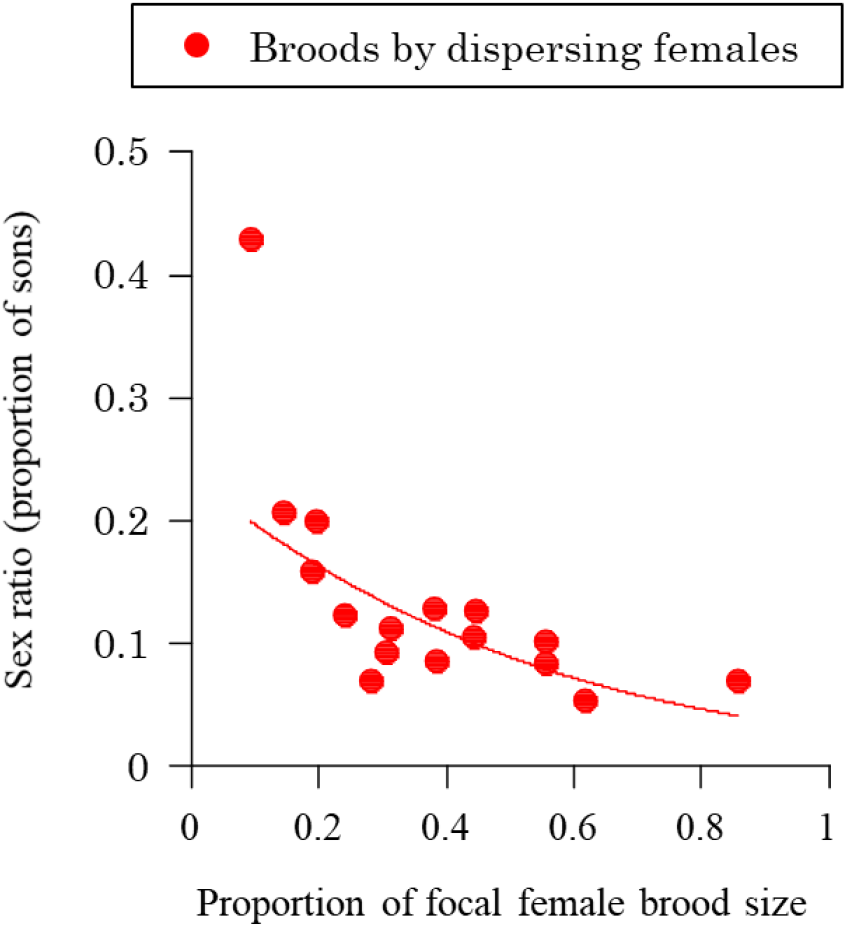
Sex ratios in response to relative brood size. Sex ratios produced by dispersing females laying eggs with other dispersers decrease with an increasing proportion of focal female brood size (focal female brood size divided by the total brood size produced), as expected in theory^39–41^. The solid red line represents the fitted line of the generalized linear mixed model assuming binomial distribution.

### Sex ratio and relatedness

Hamilton’s LMC theory can be rewritten to give the predicted sex ratio in terms of the relatedness between the offspring on a patch, rather than the number of females laying eggs on a patch^2,46^. This is useful for examining *Melittobia* sex ratios, because it can apply to cases where relatedness between offspring is influenced by multiple factors, including relatedness between mothers, and not just the number of mothers^2^.

We extended existing theory to examine the scenario where a fraction of females does not disperse, and instead remain to lay eggs with related females^28^ (Supplementary Information 3). We assumed the standard LMC scenario that *n* females lay eggs per patch, and that all mating occurs between the offspring laid in a patch. However, we also assumed that patches survive into the next generation with a probability 1 − *e*. If the patch survives, a fraction of female offspring become non-dispersers remaining on the natal patch to reproduce, and others become dispersers. In contrast, with a probability *e* the patch does not survive (goes extinct), and all female offspring become dispersers. In the next generation, extinct patches were replaced with new patches, where dispersed females from other patches can reproduce.

Our model predicts: (1) as relatedness between offspring increases, a more female-biased sex ratio is favoured; and (2) this predicted negative relationship is almost identical for dispersing and non-dispersing females (Fig. 5a and Supplementary Fig. 3-2). The relationship is almost identical because it is the increased relatedness between offspring on a patch that leads to competition and mating between relatives, and hence favours biased sex ratios^2^. Increased relatedness can come from either one or a small number of females laying eggs on a patch, or multiple related females laying eggs. For a given number of females laying eggs on a patch, non-dispersing females will be favoured to produce a more biased sex ratio (Supplementary Fig. 3 - 1), but this can also be accounted for by the extent to which their offspring will be more related (Fig. 5b). Non-dispersing females produce offspring that are more related, and so a more female-biased sex ratio is favoured (Fig. 5a). For example, if five non-dispersing females lay eggs on a patch, then the relatedness between female offspring and offspring sex ratio are both predicted to be almost the same as those for three dispersing females (Fig. 5b).

**Figure 5.**
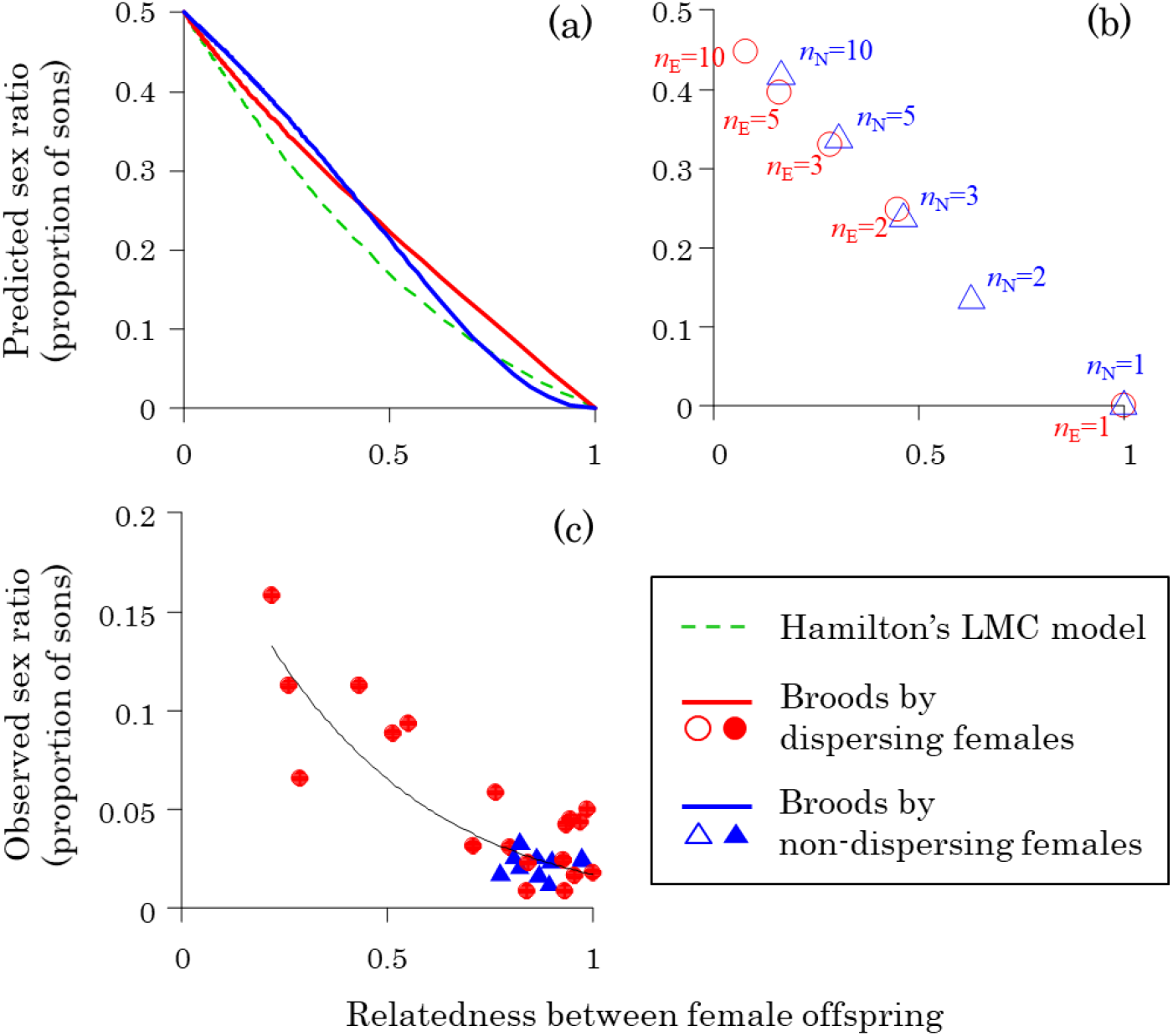
Sex ratios and relatedness. The predicted (a & b) and observed (c) relationship between the offspring sex ratio, and the relatedness between female offspring on a patch. Our model predicts that the evolutionarily stable (ES) sex ratio is negatively correlated with the relatedness between female offspring (a & b; assuming haplo-diploid genetics, and *e* = 0.66, which is assigned from the proportion of broods by dispersed females in the field). (a) The predicted relationships are almost identical for dispersed females (red line) and non-dispersed females (blue line), which are also equivalent to the prediction of Hamilton’s original local mate competition (LMC) model^2^ (green dashed line). (b) The predicted sex ratio is shown when a given number of dispersing females (*n_E_*; open red circle) or non-dispersing females (*n_N_*; open blue triangle) lay eggs on a patch. When more than one (*n* > 1) female lays eggs on a patch, non-dispersing females are predicted to exhibit more related females, and lower sex ratios. However, the overall relationship between the sex ratio and relatedness is very similar for dispersing and non-dispersing females, because the prediction for each number of dispersing females (*n_E_*) is approximately equivalent to that for a slightly lower number of non-dispersing females (*n_N_*). For example, compare the predictions for *n_E_* = 3 and *n_N_* = 5. (c) Compared across natural broods, the offspring sex ratio was negatively correlated with the relatedness between female offspring. While the sex ratios produced by dispersing females (solid circle) decrease with relatedness, the broods of non-dispersing females (solid blue triangles) were all clumped at high relatedness/ low sex ratio. The solid black line in (c) represents the fitted line of the generalized linear mixed model assuming binomial distribution for dispersing and non-dispersing females.

When we plotted the *M. australica* sex ratio data, for both dispersers and non-dispersers, there was a clear negative relationship between the sex ratio and the relatedness between female offspring (Fig. 5c; χ^2^_1_ = 25.86, *P* < 0.001). Relatedness between female offspring explained 72.2% of the variance in the offspring sex ratio. This relationship is driven by relatively continuous variation across the broods produced by dispersing females, and all the broods produced by non-dispersing females being at one end of the continuum (Fig. 5c). When more dispersing females laid eggs on a patch, this led the offspring being less related (Fig. 3a), and females produced a less female-biased sex ratio (Fig. 3b). In contrast, in the broods produced by non-dispersers, neither the relatedness between offspring or the offspring sex ratio varied significantly with the number of female laying eggs on the patch (Fig. 3). Consequently, the broods of non-dispersers were all at the extreme end of the relationship between offspring sex ratio and relatedness between female offspring (Fig. 5c).

These results show that the sex ratio behaviour of dispersing and non-dispersing females lays on the same continuum, and explains why non-dispersers do not adjust their offspring sex ratios (Fig. 5c). Non-dispersers are so related that the number of females laying eggs does not significantly influence the relatedness between their offspring (Fig. 3a). Consequently, non-dispersers always encounter extreme LMC, and are selected to produce consistently and highly female-biased offspring sex ratios (Fig. 5a).

### Can females recognize relatives?

Females could be assessing relatedness by either directly recognizing kin, or indirectly, by using whether or not they have dispersed as a cue of whether they are likely to be with non-relatives (dispersering females) or relatives (non-dispersering females)^34–37^. We conducted a laboratory experiment to test whether females directly recognize the relatedness of other females laying eggs on the same host, and adjust their sex ratio accordingly. We examined the sex ratio behaviour of females who were either: (a) laying eggs on a host alone; (b) laying eggs with a related female in the same inbred strain; (c) laying eggs with an unrelated female from a different inbred strain.

We found that there was no significant influence of relatedness between females on the offspring sex ratio that they produced. The sex ratio produced by females ovipositing with another female did not differ significantly, depending upon whether the other female was related or not (Fig. 6; χ^2^_1_ = 0.66, *P* = 0.42). As has been found previously^14,16,17^, the sex ratio produced when two females laid eggs together was slightly higher than when a female was laying eggs alone (χ^2^_1_ = 20.5, *P* < 0.001), but this shift was negligible compared to the predictions of Hamilton’s LMC theory^5^. Consequently, it appears that females cannot assess relatedness directly, consistent with previous work on *Melittobia* species^16,25^.

**Figure 6.**
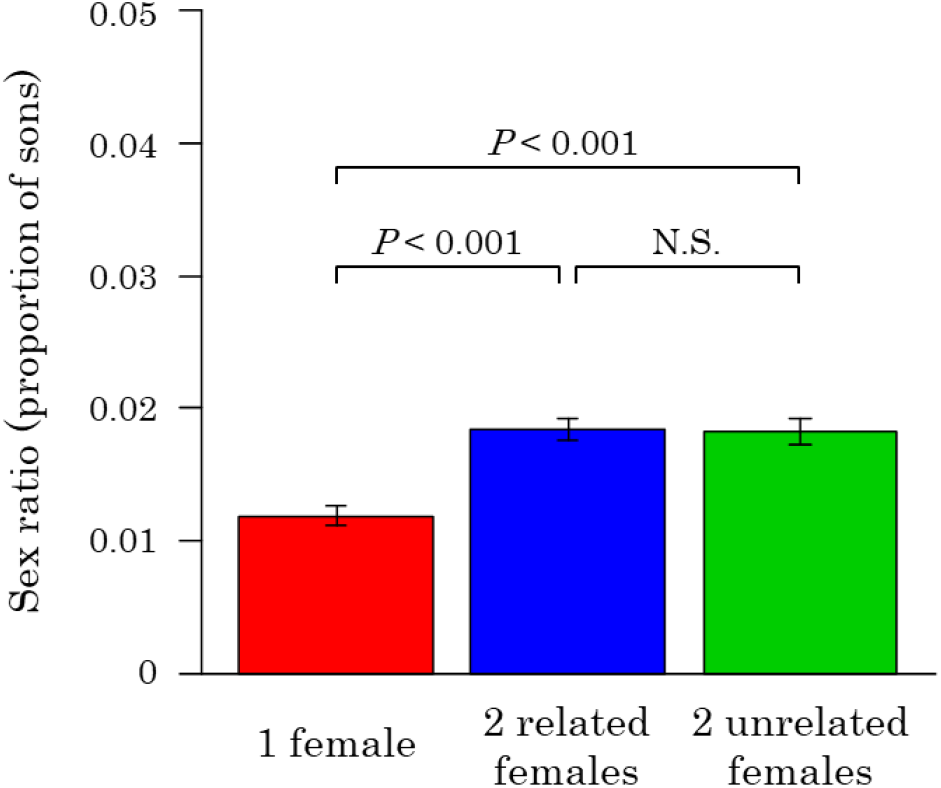
Sex ratios in the laboratory experiment. The number of replicates was 24 for each treatment. Error bars represent standard errors. Statistical values indicate the result of a pairwise comparison after correcting for multiple comparisons. Females show slightly increased sex ratios when they lay eggs with another female, as previously observed^14,16,17^. However, they do not adjust their sex ratios in response to relatedness to the females

## Discussion

We found theoretically, that when the offspring on a patch will be more closely related, females should produce more female-biased offspring sex ratios^2^ (Fig. 5a). In our model, the relatedness between offspring is determined by both how many females laid eggs on a patch, and whether those females (mothers) were had dispersed. Furthermore, we found that the predicted relationship between sex ratio and relatedness is relatively invariant to whether variation in relatedness is caused by the number of females laying eggs on a patch, or their dispersal status. Frank (1998) has previously shown that one way of thinking about and analysing LMC, is that it is the relatedness between offspring that determines the extent to which relatives are competing for mates, and mating siblings, and hence the extent of LMC^2^. Our results support Frank (1998)^2^, showing that his prediction is very little altered when allowing for variable dispersal strategies. Overall, the relatedness between female offspring was able to explain an impressive 72% of the variation in the sex ratio data, which is far higher than the average of 28% for studies on LMC^9^.

We found that, in nature, *M. australica* females adjust their offspring sex ratio in the direction predicted by our theoretical model (Fig. 5c). Females appear to do this by adjusting their offspring sex ratio in response to two factors, the number of females laying eggs on a patch, and whether they have dispersed (Fig. 3b). The influence of the number of females laying eggs has previously been demonstrated in numerous species^4^. The more females lay eggs on a patch, the less related offspring will be^2,5,38^. In contrast, the influence of dispersal status is relatively novel. Previous work on the thrip *Hoplothrips pedicularius* suggests that non-dispersing females lay more female-biased sex ratios, but it is not known if the offspring sex ratio is also adjusted in response to the number of females laying eggs^28^. What is special about the pattern in *M. australica* is that females appear to adjust their response to other females, depending upon whether they have dispersed or not – only dispersed females lay less female-biased sex ratios when more females are laying eggs on a patch (Fig. 3b). Consequently, *M. australica* females appear to use a relatively sophisticated strategy to adjust their offspring sex ratios in response to the extent of LMC, as measured by the relatedness between the offspring that will develop on that patch (Fig. 5).

*M. australica* females appear to assess their relatedness to other females indirectly, by their dispersal status, rather than directly, by genetic cues. The optimal sex ratio depends upon relatedness to other females laying eggs on the same patch, and so females will be selected to assess relatedness. Our laboratory experiment suggests that females cannot assess relatedness directly (Fig. 6). The direct assessment of kin via genetic cues appears to be rare in parasitoids, with most studies finding that females do not use genetic cues to assess relatedness to other females^16,25,35–37,47^. An exception is provided by the bethylid *Goniozus nephantidis*, where females appear to be able to assess relatedness directly, and adjust their offspring sex ratio accordingly^34,36^. A possible reason for the rarity of direct genetic kin recognition is that it is not evolutionarily stable, because selection will favour more common markers, and hence eliminate the genetic variability required for kin recognition^48–50^. In contrast, dispersal status appears to be an excellent indirect (environmental) cue of relatedness^28–30^ – when females do not disperse, they are closely related to the other females laying eggs on a patch, leading to their offspring being highly related (Fig. 3a).

While the pattern of sex ratio adjustment in *M. australica* is in the direction predicted by theory, the fit to theory is qualitative not quantitative. The offspring sex ratios are more female biased than predicted by theory (Fig. 5). One possible explanation for that is a cooperative interaction between related females^6,51–53^. In *Melittobia*, females favor ovipositing on hosts parasitized by other females rather than intact hosts (J. A. unpublished data). In addition, a larger number of *Melittobia* females are likely to be advantageous to cooperatively make tunnels in host nests, and to fight against mite species that lives symbiotically with host species^54,55^. Cooperative interactions have previously been suggested to favour an increased proportion of female offspring in a range of organisms, including other parasitoids, bees, beetles, and birds^37,51,56–60^. A complication here is that although limited dispersal increases relatedness between encountering individuals, it can also increase competition between the related individuals, and so reduce selection for female-biased sex ratios^32,33,46,53,61^. However, in the case of *Melittobia* species, overlapping generations, inelasticity, dispersing with relatives, and open sites could negate this increased competition^33,62–65^ (Supplementary Information 3)

To conclude, our results provide an explanation for the *Melittobia* sex ratio scandal. Laboratory experiments on four *Melittobia* species have found that females produce lay extremely female-biased sex ratios (1 – 5% males), that change little with increasing number of females laying eggs on a patch^14,18–20,22^ (Fig. 1). In laboratory experiments, females were not able to disperse, and so they were likely to have behaved as non-dispersers, who would normally be on a patch with close relatives. We found that, in natural conditions, non-dispersers behave as if other females are close relatives, and show little response to the number of females laying eggs on the same patch (Fig. 3b). So rather than being scandalous, the sex ratio behavior of *Melittobia* appears to reflect a refined strategy, that also takes account of indirect cues of relatedness.

## Methods

### Parasitoid wasps

*Melittobia australica* (Hymenoptera: Eulophidae) is a gregarious parasitoid wasp that mainly parasitizes prepupae of various solitary wasp and bee species nesting above ground. The host species build nests containing brood cells, in which offspring develop separately while eating food provided by their mothers. Several generations (5 or more on the main island of Japan) appear per year in *Melittobia* species, while most of the host species are mainly univoltine. Multiple generations of *Melittobia* may occur in a single host patch. Therefore, females laying eggs in *Melittobia* could be derived from other broods in the same host patch or disperse from other patches (Fig. 2). Once a female finds a host, she continues to lay eggs on the surface of the host, and is potentially able to produce several hundred eggs on a single host as long as the host resources are sufficient. Similar to other hymenopteran species, *Melittobia* exhibits a haplo-diploid sex determination, in which male and female individuals develop from unfertilized and fertilized eggs, respectively. Hatched larvae develop by sucking host haemolymph from the outside, and offspring sequentially emerge from previously laid eggs. The adults of males are brachypterous, and do not disperse from their natal host cells, in which they mate with females that develop on the same host^21^.

### Field sampling

To collect the *Melittobia* broods from the field, we used bamboo traps, in which the host species build nests to develop their offspring. We set up 7 – 32 bamboo traps at the end of July or the beginning of August from 2011 – 2019 within an area with a 2 km radius in and around the campus of Kanagawa University in Hiratsuka, Kanagawa, Japan (Supplementary Table 2-1). The traps were horizontally fixed to the trunks or branches of trees at approximately 1.5 m above the ground. Each bamboo trap consisted of a bundle of 20 dried bamboo canes. Each cane (200 – 300 mm long and 10 – 15 mm internal diameter) was closed with a nodal septum at one end and open at the other end. The traps were collected between December and the following March, when the individuals of the host wasps and bees in the cells were undergoing diapause at the prepupal stage. The traps were brought back to the laboratory, where all of the canes were opened to inspect the interior. Along with the traps that were set up in summer as described above, we conducted field sampling in the spring of 2015, 2017, and 2018. Bamboo canes containing nests with host prepupae that were collected in winter and kept in a refrigerator were exposed in the field for 1 – 2 months.

In the laboratory, all individuals of the solitary wasps and bees and their kleptoparasites and parasitoids in the bamboo canes were counted and identified at the species level according to the morphology of the juveniles and the construction of the nests (Supplementary Table 2-2). If they were parasitized by *Melittobia* species, we recorded the host and *Melittobia* species (Supplementary Table 2-3). The adults and the dead bodies of *Melittobia* females in the host cells or cocoons could be regarded as mothers that laid eggs in the broods, if the next generation had not yet emerged. We could distinguish female offspring from mothers based on the filled abdomen of emerged females. We collected the mothers, and preserved them in 99.5% ethanol. After removing the mothers, the parasitized hosts were individually incubated at 25°C until the offspring emerged. Emerged individuals were counted, sexed, and preserved in 99.5% ethanol.

### Molecular analysis

For the analysis of sex ratios and molecular genetics, we only adopted the broods of *M. australica* among the two collected species (Supplementary Table 2-4), because this species was dominant in the present sampling area, and most of the applied microsatellite markers indicated below were only valid for this species^27^. We also only included broods in which all the individuals of the offspring generation had not yet started to emerge. The existence of emerged offspring in the host cells may indicate that some females had already dispersed. Finally, we excluded clutches in which 16 or fewer individuals emerged, or the majority of juveniles were destroyed by an accidental event such as host haemolymph flooding. For genotyping, we analysed 16 female offspring that were randomly selected from all emerged females in each clutch. For mothers and male offspring, we genotyped all individuals if the number of individuals was less than 16; otherwise, 16 randomly selected individuals were analysed.

We used the boiling method for DNA isolation (Abe et al. 2009). DNA was isolated from the whole body for male individuals, or from the head and thorax for female individuals to prevent the contamination of spermatozoa from their mates. For the genomic analysis, we selected 16 microsatellite markers out of 19 markers^27^ to avoid the use of markers that might show genetic linkage with other loci. The detailed polymerase chain reaction methods are described elsewhere^27^. The microsatellite genotypes of each individual were identified from the amplified DNA fragments through fragment analysis with an ABI 3130 capillary sequencer (Life Technologies, Carlsbad, CA, USA) and Peak Scanner software version 1.0 (Life Technologies, Carlsbad, CA, USA). Based on the genotyping data, the average relatedness between the individuals on the broods and inbreeding coefficient were calculated using the software Relatedness version 4.2b^66^. The broods were equally weighted in all analyses. We estimated the relatedness between the female offspring in a brood as a representative metric of patch relatedness, because we could obtain a sufficient number of female offspring from all the patches.

Although adult females that were captured in the broods of developing offspring could be regarded as females laying eggs, all of the females might not necessarily contribute the broods, and other females might depart before collection. We conducted a parentage analysis to reconstruct the genotypes of females laying eggs (mothers) from the genotypic data of the offspring following the Mendelian rules under haplo-diploidy (Supplementary Table 2-6). We first assigned the minimum number of captured adult females as the candidate mothers. If we could not identify the candidates from the captured females, we assumed that additional females produced offspring in the brood, and rebuilt their genotypes. For assignment, we adopted the solution with the minimum number of mothers producing broods. We also assumed that mothers mated with multiple but possibly minimal numbers of males. Consequently, the genotypes of the mothers could be uniquely determined for dispersers, and the number of mothers was strongly correlated with that of captured females (Pearson’s correlation: *r* = 0.84, *t* = 6.51, *df* = 17, *P* < 0.001). However, we could not uniquely determine the genotypes of mothers for non-dispersers, because of the high similarity of the genotypes of the captured females. Then, we used the number of captured females as the number of mothers for non-dispersers, although this may be an overestimation.

### Laboratory experiment

A female was introduced into a plastic case (86 mm in diameter and 20 mm in height) (a) alone, (b) with a related female from the same inbred strain, or (c) with an unrelated female from a different strain, and allowed to lay eggs on a prepupa of *Chalicodoma sculpturalis* for 12 days. All the females of the same strains were collected from a single prepula of *C. sculpturalis* that was parasitized by 8 females randomly chosen among the laboratory strains. The laboratory strains were the Amami strain (AM) and the Shonan 1 or 2 strain (SN1 or SN2), which were initiated by wild-caught wasps from single hosts collected from Amami Oshima Island in 2007 and from Hiratsuka in 2017, respectively. Amami Oshima and Hiratsuka are located approsimately 1200 km away, and are separated by sea. The Shonan 1 and 2 strains originated in the same population, but were developed from individuals caught by different bamboo traps. The Amami and Shonan strains were maintained in the laboratory for approximately 70 and 10 generations, respectively, during which several dozen females from previous generations were allowed to lay eggs on prepupae of *C. sculpturalis* to produce the subsequent generations. This procedure mimics the situation in which non-dispersed females produce a new brood on another host in the same patch in the field. Before the experiment, a female was allowed to mate with a male from the same strain in the manner described elsewhere^17^. The emerged offspring were counted and sexed. In the treatment involving two females from different strains, all male and 16 randomly collected female offspring were genotyped for one of the microsatellite loci^27^, and the offspring sex ratios of the two ovipositing females were estimated separately using the ratio of the genotyped individuals. In the treatment involving females from the same strain, the two females were assumed to equally contribute to the clutches. The experiment was conducted at a temperature of 25°C and a photoperiod of L16:D8 h.

### Statistical analysis

We analysed all data using generalized linear mixed models implemented in statistical software R version 3.6.1 ^67^. The data on the number of females laying eggs or offspring, the sex ratio, the injury frequency, and relatedness were fitted with Poisson, binomial, binomial, and beta distributions, respectively. Because, multiple broods in the same bamboo canes were estimated to be produced by females from the same hosts in some broods by non-dispersers, we added the bamboo cane to the models as a random effect. In addition, we added the individual brood as a random effect for count data (the number of females laying eggs or offspring) or proportional data (sex ratio) to resolve the problem of over-dispersion. We first ran full models including all fixed effects of interest and random effects, and then simplified the models by removing non-significant terms (α > 0.05) from the least significant ones to arrive at the minimum adequate models^68^. The statistical significance was evaluated by using the likelihood ratio test comparing the change in deviance between the models. In multiple comparisons, significance thresholds were adjusted using the false discovery rate^69^.

### Theoretical model

We assume a population consisting of infinite discrete patches, in which emerged male and female offspring mate at random^70^ (Wright’s islands model of dispersal). We define the patch extinction rate *e* as the probability that patches will go extinct, and be replaced by new patches. If the patches go extinct (which occurs with a probability of *e*), all developed female offspring on the patches disperse to other patches. In contrast, if a patch has survived (with a probability of 1 – *e*), female offspring do not disperse with a probability of 1 – *d*, or otherwise disperse with a probability of *d*. Non-dispersers remain on their natal patch, in which *n* randomly selected females reproduce in the next generation. Dispersers from all the patches compete for reproduction on the newly created patches, in which *n* randomly selected females reproduce. We use the subscript *i* to denote dispersers (*i* = E) or non-dispersers (*i* = N). Let *x_i_* denote the sex ratio of a focal female, *y_i_* denote the average sex ratio in the same patch as the focal female, and *z_i_* denote the average sex ratio in the whole population. Then, the relative fitness of a daughter of the focal female 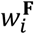 and the relative fitness of a son of the focal female 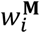 may be written as follows:

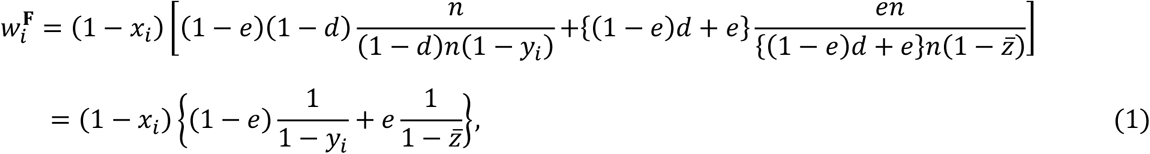

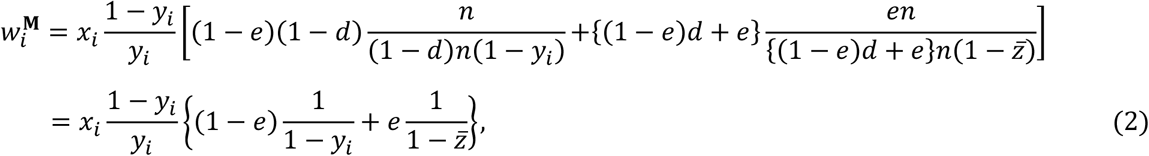

respectively, where 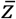 represents the metapopulation wide average sex ratio:

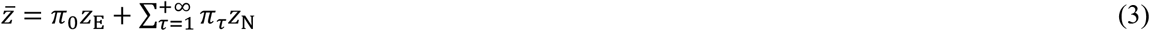

and *π_τ_* is the frequency of patches that were recolonized during *τ* generations. Note that the parameter *d* is cancelled out as dispersers compete only with dispersers, while non-dispersers only with non-dispersers. By applying the neighbour-modulated fitness approach to kin selection methodology^2,71–74^, we find that an increased sex ratio is favoured by natural selection for dispersers, if:

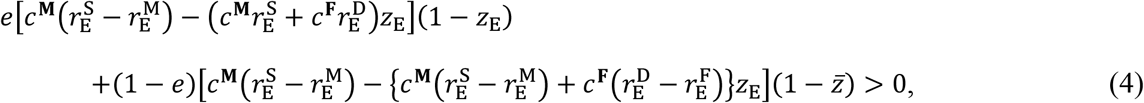

and for non-dispersers, if:

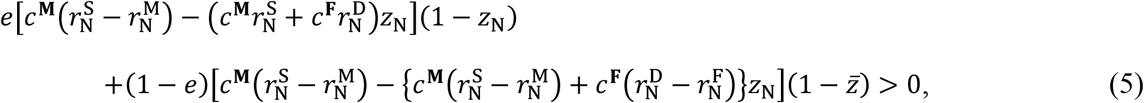

where *c*^**M**^ and *c*^**F**^ are the class reproductive values for females and males, respectively, 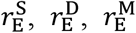 and 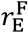 are the kin selection coefficients of relatedness for a daughter, a son, a random male offspring on the same patch, and a random female offspring on the same patch, respectively, from a perspective of a disperser, and 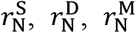 and 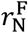 are the average kin selection coefficients of relatedness for a daughter, a son, a random male offspring on the same patch, and a random female offspring on the same patch, respectively, from a perspective of a non-disperser. Substituting the terms of reproductive values and relatedness, we may derive the convergence stable sex ratios for dispersers 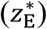 and non-dispersers 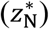. If we replace *e* with 1 − (1 − *d*)^2^, where *d* is female dispersal rate after mating, we recover Gandner et al.’s (2009) results^33^. A full derivation is given in Supplementary Information 3.

## Acknowledgements

We thank Mahiro Abe and Ryo Abe for field assistance. Funding was provided by a Japan Society for the Promotion of Science grant-in-aid for scientific research (JSPS KAKENHI grant 17K07574) to J. A.

## Author contributions

J. A., Y. K., and S. A. W. initiated, planned, and coordinated the project; J. A. collected and analysed field, molecular, and laboratory data; R. I. and J. A. constructed the theoretical model; K. T. conducted the genetical analyses; and all authors wrote the paper.

## Supplementary information 1

### Field data

We obtained 46 broods of *Melittobia australica* with bamboo traps placed in the wild, and 11 broods by reexposing upparasitised hosts in the field (Supplementary Table 2-1, 2-2). All of the emerging offspring were obtained with sufficient confidence from 29 of the collected broods, so we analysed these broods (Supplementary Table 2-3). Our parentage analysis with the microsatellite genotypic data indicated that all but two of the 29 broods were founded by either only dispersers or only non-dispersers (Supplementary Table 2-3, S2-4). In the two other broods, non-dispersers produced male and female offspring, while a single disperser added an all-male clutch. In both broods, 5 non-dispersers were collected with developing offspring on the host, while the disperser was not collected (Supplementary Table 2-4), suggesting that she had left the host before collection. The single dispersers produced 75.0% (50.3/67) and 87.5% (66.5/76) male individuals, respectively, in the broods. We were not certain whether the females producing all-male clutches were virgins, although their behaviour is different from that of virgin females in the laboratory, in which they produced only a few (maximally 9) male offspring before they mate with their own sons (Abe et al. 2010).

### Genetic structure

To examine genetic differentiation depending on the behaviour of females laying eggs, hierarchical population structure was analysed in microsatellite data using the R package *hierfstat* (Goudet 2005). When we consider a structure, in which the entire population (population) is divided into two groups of dispersers and non-dispersers (group), and each group is divided by the genetic lineages of the females (lineage), the effect of group was not significant (*F*_group/population_ = 0.011, *P* = 0.43), suggesting that there is no genetic differentiation between non-dispersers and dispersers. When we instead divided the entire population into two groups of all-male producers and the other females, the effect of group was not significant (*F*group/population = –0.020, *P* = 0.30), suggesting that there was no genetic differentiation between the females that produced all-male clutches and those that produced clutches containing both sexes.

### The number of females laying eggs

Over 10-fold more non-dispersers laid eggs on a single host (mean ± SD = 16.6 ± 11.9) compared with dispersers (mean ± SD = 1.4 ± 1.1; Supplementary Table 2-3; χ^2^_1_ = 16.54, *P* < 0.001). The effect of host species on the number of females laying eggs per brood was marginally non-significant (χ^2^_5_ = 10.81, *P* = 0.055).

### Brood size

Non-dispersers produced more offspring than dispersers in a single brood, although there were no significant effects of females laying eggs or host species (Supplementary Table 1-1a, 2-3). However, the number of females laying eggs was highly related to the dispersal status of the females, as shown above. When we analysed the model after removing the dispersal status term, offspring number significantly increased with the number of females, but host species was still non-significant (Supplementary Table 1-1b).

**Supplementary Table 1-1a.**
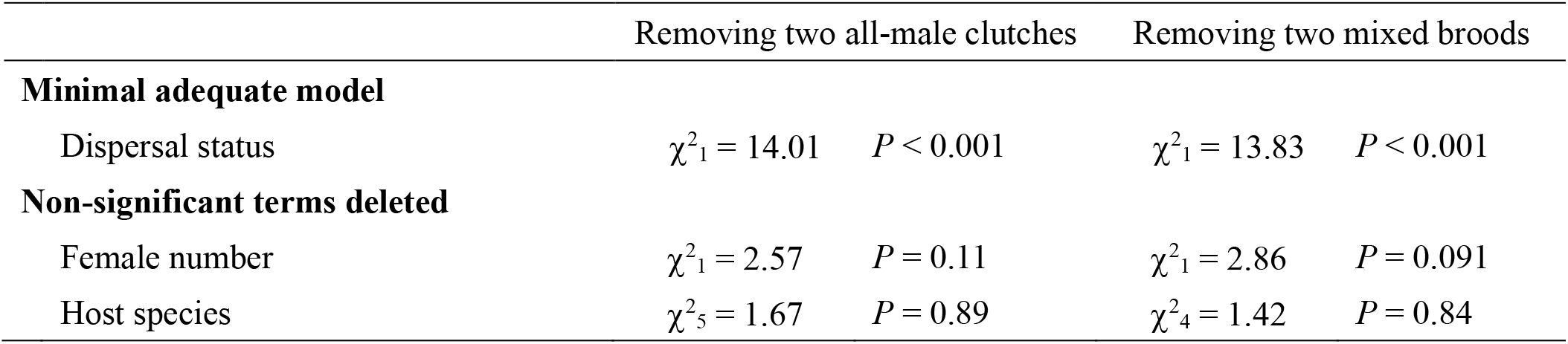
Analysis of total brood size (with dispersal status).

**Supplementary Table 1-1b.** Analysis of total brood size (without dispersal status).

|  | Removing two all-male clutches |  | Removing two mixed broods |  |
| --- | --- | --- | --- | --- |
| **Minimal adequate model** |  |  |  |  |
| Female number | $\chi^2_1 = 9.61$ | $P = 0.002$ | $\chi^2_1 = 13.52$ | $P < 0.001$ |
| **Non-significant terms deleted** |  |  |  |  |
| Host species | $\chi^2_5 = 3.06$ | $P = 0.69$ | $\chi^2_4 = 2.13$ | $P = 0.71$ |

### Relatedness

We adopted relatedness between female offspring in a brood to assess the kinship between individuals on a patch, because we could obtain a sufficient number of female offspring in all the broods analysed (Supplementary Table 2-3). Relatedness between female offspring showed a significant interaction between the number of females laying eggs and the dispersal status of the females (Fig. 3a), although brood size and host species were non-significant (Supplementary Table 1-2a). When we analysed the model for each dispersal status separately, relatedness significantly decreased with an increasing female number in the broods of dispersers (Supplementary Table 1-2b), but relatedness was independent of female number in the broods of non-dispersers (Supplementary Table 1-2c).

**Supplementary Table 1-2a.** Analysis of relatedness between female offspring in a brood (including both type of females).

|  | Removing two all-male clutches |  | Removing two mixed broods |  |
| --- | --- | --- | --- | --- |
| **Minimal adequate model** |  |  |  |  |
| Female number | NA |  | NA |  |
| Dispersal status | NA |  | NA |  |
| Female number : Dispersal status | $\chi^2_1 = 12.34$ | $P < 0.001$ | $\chi^2_1 = 11.97$ | $P < 0.001$ |
| **Non-significant terms deleted** |  |  |  |  |
| Brood size | $\chi^2_1 = 2.67$ | $P = 0.10$ | $\chi^2_1 = 2.31$ | $P = 0.13$ |
| Host species | $\chi^2_5 = 5.47$ | $P = 0.36$ | $\chi^2_4 = 5.09$ | $P = 0.28$ |

**Supplementary Table 1-2b.** Analysis of relatedness between female offspring in a brood (only dispersers).

|  | Removing two all-male clutches |  | Removing two mixed broods |  |
| --- | --- | --- | --- | --- |
| **Minimal adequate model** |  |  |  |  |
| Female number | $\chi^2_1 = 10.15$ | $P = 0.001$ | $\chi^2_1 = 10.15$ | $P = 0.001$ |

**Supplementary Table 1-2c.**
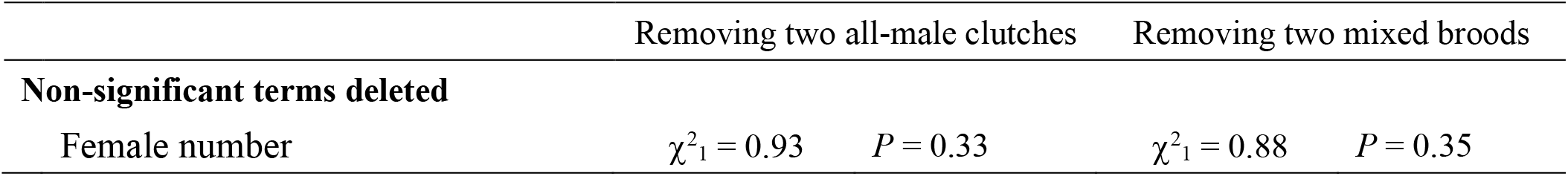
Analysis of relatedness between female offspring in a brood (only non dispersers).

### Sex ratio

Sex ratios were clearly categorized into two groups depending on the dispersal status of females (Fig. 3b): the interaction term between the number of females laying eggs and their dispersal status was significant (χ^2^_1_ = 18.95, *P* < 0.001), although host species and brood size were not significant (Supplementary Table 3-1a). Separate model analysis for each dispersal status showed that dispersers increased the sex ratio by increasing the number of females laying eggs (Supplementary Table 3-1b), whereas non-dispersers did not (Supplementary Table 3-1c). When we analysed sex ratios against relatedness between female offspring incorporating the broods of both dispersers and non-dispersers, a significant negative relationship between the sex ratio and relatedness was found (Supplementary Table 1-4).

**Supplementary Table 1-3a.** Analysis of sex ratio (both type of females).

|  | Removing two all-male clutches |  | Removing two mixed broods |  |
| --- | --- | --- | --- | --- |
| **Minimal adequate model** |  |  |  |  |
| Female number | NA |  | NA |  |
| Dispersal status | NA |  | NA |  |
| Female number : Dispersal status | $\chi^2_1 = 18.69$ | $P < 0.001$ | $\chi^2_1 = 18.70$ | $P < 0.001$ |
| **Non-significant terms deleted** |  |  |  |  |
| Host species | $\chi^2_5 = 8.40$ | $P = 0.16$ | $\chi^2_4 = 7.40$ | $P = 0.12$ |
| Brood size | $\chi^2_1 = 0.095$ | $P = 0.76$ | $\chi^2_1 = 1.60$ | $P = 0.21$ |

**Supplementary Table 1-3b.** Analysis of sex ratio (only dispersers).

|  | Removing two all-male clutches |  | Removing two mixed broods |  |
| --- | --- | --- | --- | --- |
| **Minimal adequate model** |  |  |  |  |
| Female number | $\chi^2_1 = 14.62$ | $P < 0.001$ | $\chi^2_1 = 14.62$ | $P < 0.001$ |

**Supplementary Table 1-3c.** Analysis of sex ratio (only non dispersers).

|  | Removing two all-male clutches |  | Removing two mixed broods |  |
| --- | --- | --- | --- | --- |
| Non-significant terms deleted |  |  |  |  |
| Female number | $\chi^2_1 = 0.56$ | $P = 0.46$ | $\chi^2_1 = 0.015$ | $P = 0.90$ |

**Supplementary Table 1-4.** Analysis of sex ratio against relatedness between female offspring in a brood.

|  | Removing two all-male clutches |  | Removing two mixed broods |  |
| --- | --- | --- | --- | --- |
| **Minimal adequate model** |  |  |  |  |
| relatedness | $\chi^2_1 = 25.86$ | $P < 0.001$ | $\chi^2_1 = 24.27$ | $P < 0.001$ |

## Laboratory data

### Brood size

The number of offspring produced by a female was not influenced by the treatment (Supplementary Fig. 1-1a; χ^2^_2_ = 0.70, *P* = 0.70) or strain (χ^2^_2_ = 3.84, *P* = 0.15).

### Sex ratio

Although the offspring sex ratio was significantly different depending on the treatment (χ^2^_2_ = 21.27, *P* < 0.001), this relied on the difference in foundress numbers (χ^2^_1_ = 20.5, *P* < 0.001), but the sex ratios produced by two related and unrelated females were not significantly different (Fig. 5; χ^2^ = 0.66, *P* = 0.42). The strain did not have a significant effect on the sex ratio (Supplementary Fig. 1-2b; χ^2^ = 1.50, *P* = 0.47).

### Injury level

Fortuitously, we observed fighting between females in the experiment, which has rarely been documented in *Melittobia* (Matthews & Deyrup 2007). Parts of the antennae and legs of females were likely to be cut off by the opponent female during the 8 days after the introduction of the females. However, the frequency of the injured females was not influenced by relatedness (Supplementary Fig. 1-1c; χ^2^_1_ = 0.05, *P* = 0.82), although female pugnacity significantly varied among the strains of females (χ^2^_1_ = 18.32, *P* < 0.001). Ultimately, we found no evidence that females adjust their behaviour depending on relatedness. Moreover, females could potentially assessed relatedness indirectly on the basis of environmental cues, such as recognizing whether the opponent females emerged from the same or different host. However, the present experiment, in which all females of the same strain that were used were developed on the same host, suggested that this possibility is not the case in the studied species.

**Figure S1-1.**
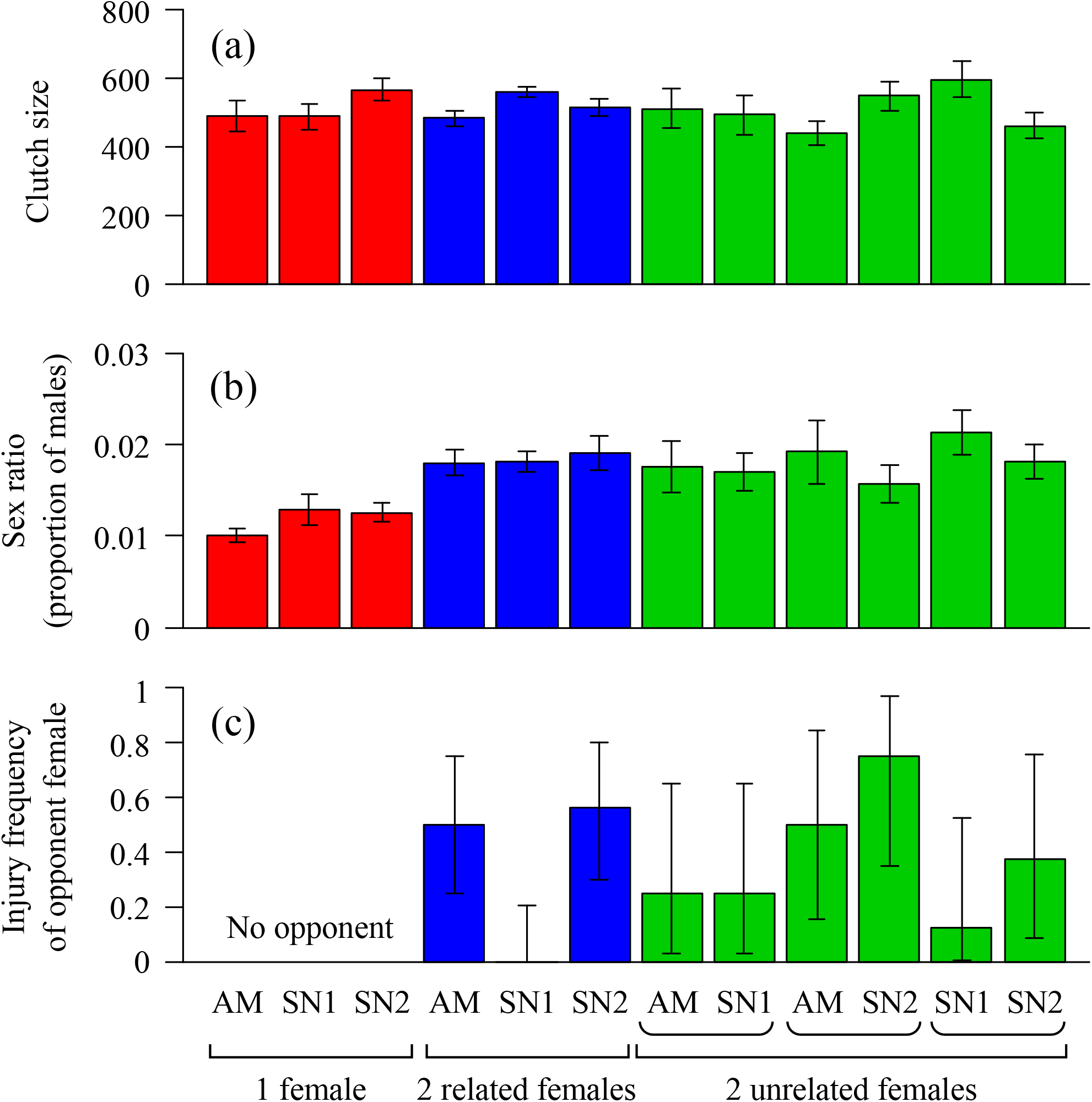
Clutch size (a), sex ratio (b), and injury frequency of opponent female (c) depending on treatment
regulating female number and their relatedness, and the strains of the females. Error bars represent standard errors (a, b) and 95% binomial confidence intervals (c). The number of replicates was 8 for each strain in all the treatments

## 1 Model assumptions

We use a spatially implicit model of dispersal (Wright’s islands model; Wright 1931), in which each patch may go extinct at a probability *e*. If patches go extinct, the same number of empty patches are recolonized in the next generation. Therefore, each patch is characterized by the age *τ*, where *τ* is the number of generations that have passed since a patch was recolonized.

## 2 Patch dynamics

Let *π_τ_* be the frequency of the patches aged *τ*. Under completely random extinction, the frequency of patch ages is updated by:

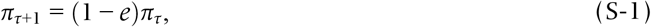

with

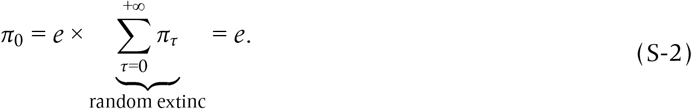

Hence, the stationary distribution of the patches aged *τ* is given by:

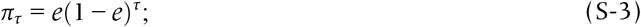

that is, *τ* follows a geometric distribution.

## 3 Consanguinity

Under the assumptions that females and males mate within their natal patches and *n* non-dispersing females reproduce in persistent patches, while *n* dispersing females from other patches reproduce in recolonized patches, consanguinity coefficients (the probability that randomly taken homologous genes of interest are identical by descent) may be derived for diploid and haplodiploid populations as below.

### 3.1. Diploidy

The consanguinity coefficient *f_τ_* between a random mating pair on a patch aged *τ* is given by a well known recursion (Taylor 1988a,b; Frank 1998; Rousset 2004; Lehmann 2007; Gardner et al. 2009):

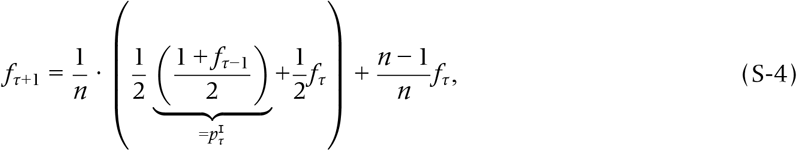

in which identical by decent (IBD) between mating partners occurs when (i) they share share the same mother 1 ∕ *n*, in which case the consanguinity is given by (i-a) the probability that both genes are from the same-sex parent 1/2 (i.e., both from the mother, or both from their father) times the consanguinity to self 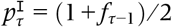 in the previous generation plus (i-b) the probability that their genes derive from opposite-sex parents 1/2 (i.e., one from mother and the other from father) times the consanguinity between the parents in the previous generation (*f_τ_*), or when (ii) they have different mothers (1 − 1 / *n*), in which case the consanguinity is given by the probability that two distinct adults share the common ancestor in the previous generation *f_τ_*.

The “initial” consanguinity (gifted by dispersers) reads:

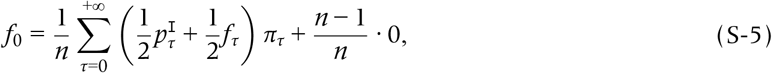

which is reasoned as follows: with a probability 1/*n*, two female offspring share the dispersed mother, in which case their consanguinity is the metapopulation-wide average of (1 + 3*f_τ_*_−1_) / 4. With a probability of 1 − 1 / *n*, two female offspring have different, dispersed mothers, in which case consanguinity is null.

Also we need to construct a recursion for 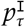, which for *τ* ≥ 1 reads:

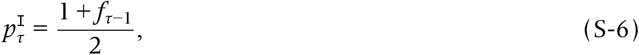

because with probability 1/2, the same homologous allele is sampled, in which case IBD is 1, and with probability 1/2, the other is sampled, in which case IBD is given by the consanguinity with her mating partner *f_τ_*_−1_. The initial condition for 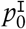 is given by:

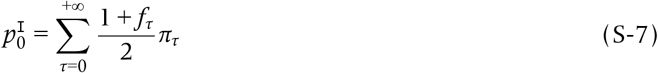

From these, we get the consanguinity coefficient between a random adult female and her own sons or daughters (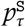 or 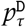, respectively) and that between a random adult female and a random offspring born in the same patch (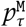 or 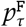, respectively) for *τ* ≥ 0:

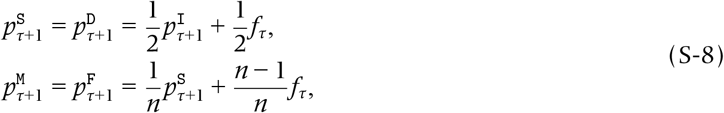

the first line of which reads as the probability that offspring’s allele derives from mother (1/2) times the probability that this allele is IBD with the mother 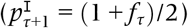, plus the probability that the allele derives from father (1/2) times the probability that this allele is IBD with the mother (*f_τ_*). The second line is because, for a given allele sampled from a random adult female, an allele sampled from one of the offspring born in the same patch derives from the adult female (1 / *n*; in which case the consanguinity is 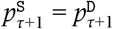) or another adult female (1 − 1 / *n*; in which case the consanguinity is *f_τ_*).

The initial conditions for *τ* = 0 (with Eqn (S-5)) are given by:

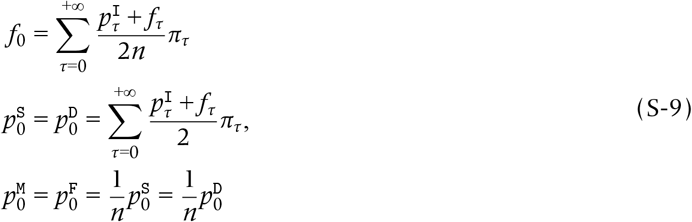

where the first line follows because the consanguinity of a mother to one of her own offspring is the arithmetic mean (1/2) for herself (the former) and her mate (latter).

### 3.2. Haplodiploidy

We denote the consanguinity of mating partners on the patch aged *τ* by *f_τ_*; the average consanguinity of two female offspring sharing the same patch aged *τ* by *ϕ_τ_*; the average consanguinity between two male offspring sharing the same patch aged *τ* by *μ_τ_*.

The consanguinity between a pair of offspring male and offspring female on the same patch aged *τ* +1 is given by the probability that they share the same mother (1/*n*) times the consanguinity of full sibs (with probability 1/2, the offspring female derives her gene from her mother as does the offspring male, in which case the consanguinity is 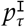; with probability 1 /2, the offspring female derives her gene from the father while the offspring male derives his gene from his mother, in which case the consanguinity is *f_τ_*) plus the probability that they do not share the same mother (1 − 1 / *n*) times the probability that their mothers are both non-disperser (which is 1 since *τ* + 1 ≥ 1) times the consanguinity of the parents by which their genes are transmitted (which is 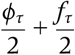). That is:

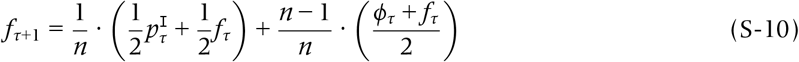

(Taylor 1988a,b; Frank 1998; Rousset 2004; Lehmann 2007; Gardner et al. 2009).

The average consanguinity of two female offspring sharing the same patch aged *τ* > 0, *ϕ_τ_*, is given by the probability that they share the same mother (1 / *n*) times the consanguinity of full sisters 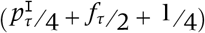 plus the probability that they do not share the same mother (1 − 1/ *n*) times the probability that their mothers are both non-dispersers (which is 1 since *τ* + 1 ≥ 1) (with probability 1 / 4 they both derived their genes from their mothers, in which case the consanguinity is *ϕ_τ_*; with probability 1 /2 they derived their genes from opposite-sex parents, in which the consanguinity is *f_τ_*; with probability 1 /4 they both derived their genes from their fathers, in which case the consanguinity is *μ_τ_*). That is,

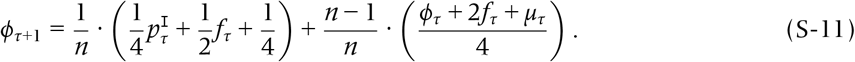

The average consanguinity of two male offspring sharing the same patch (aged *τ* + 1 ≥ 1), *μ_τ_*, is the probability that they share the same mother (1 /*n*) times the probability of full brothers 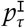 plus the probability that they do not share the common mother 1 − 1 / *n* times the probability that both of their mothers are non-disperser (which is 1 since *τ* + 1 ≥ 1) times the average consanguinity of two female offspring on the same patch *ϕ_τ_*. That is:

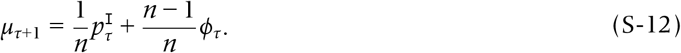

Finally, 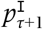(the consanguinity for herself) follows a recursion given by:

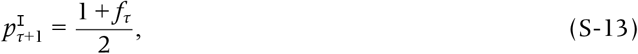

as with probability 1/2, the same allele is sampled twice (in which case IBD is 1) and with probability 1/2, two different homologous alleles are sampled (in which case IBD is given by the inbreeding coefficient in the previous generation, *f_τ_*).

For a patch that is newly recolonized (i.e., aged *τ* = 0), kinship is possible for sibs (i.e., only by sharing the mother). The consanguinity between a pair of mating partners on the patch is given by the probability that they share the mother (1 /*n*) times the spatial average of the consanguinity of full sibs:

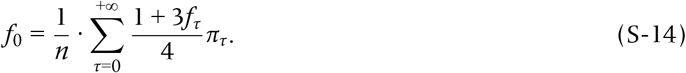

Similarly,

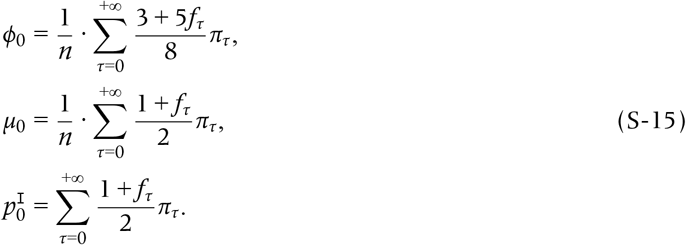

(*f*_0_, *ϕ*_0_, *μ*_0_, 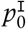) gives the initial condition of the recursion. Solved recursively, (*f_τ_*, *ϕ_τ_*, *μ_τ_*, 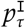) specifies the consanguinity between a mating pair, female offspring, and male offspring, respectively. From these, we get:

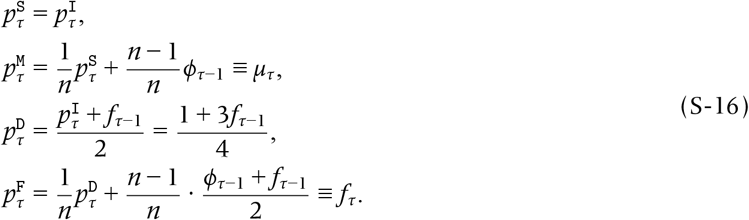

The initial condition for *τ* = 0 is given by:

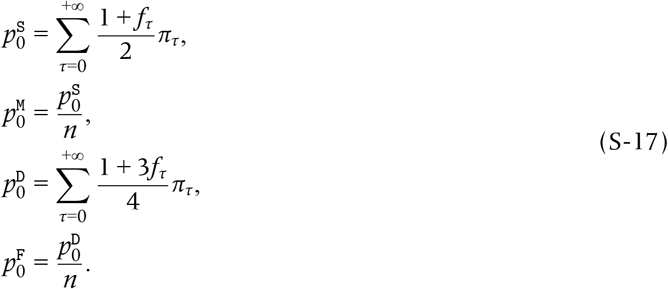

## 4 Average and initial consanguinity coefficients

### 4.1. Diploidy

We consider the average values of *f_τ_* and 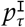 over the distribution *π_τ_*:

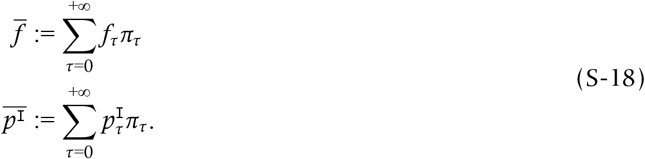

By multiplying *π_τ_*_+1_ = (1 − *e*)*π_τ_* with Eqns (S-4) and (S-6) and then summing up both sides over *τ* = 0 to ∞, we get:

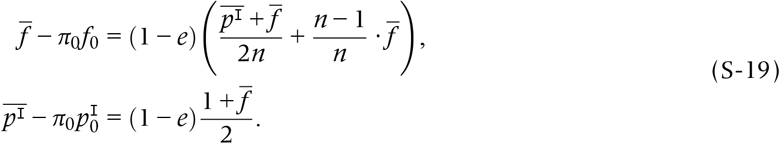

With Eqn (S-7), we get:

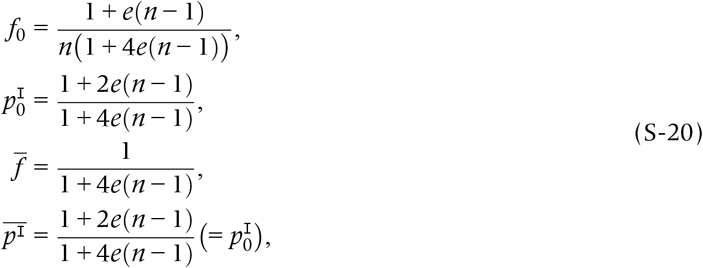

which recovers Gardner *et al*.’s (2009) results by replacing *e* with 1− (1 − *d*)^2^ (where *d* is female-dispersal rate after mating). From this calculation, one may see that the well-known recursions for the consanguinity coefficients (Taylor 1988a,b; Frank 1998; Rousset 2004; Lehmann 2007; Gardner et al. 2009) are evaluated at metapopulation-wide average over the patch-age distribution.

### 4.2. Haplodiploidy

Similarly, let us denote the spatially averaged *f_τ_*, *ϕ_τ_*, *μ_τ_* over the distribution *π_τ_* by 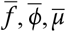, repsectively:

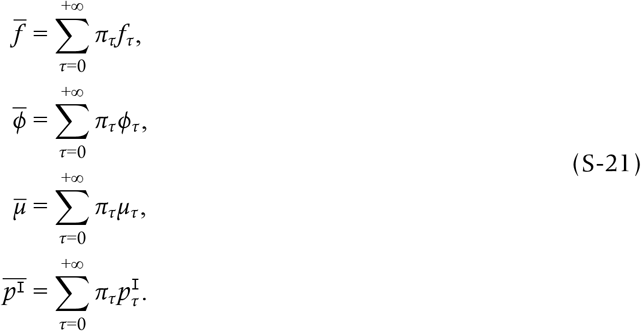

Then the initial values (*f*_0_, *ϕ*_0_, *μ*_0_, 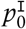) are written as

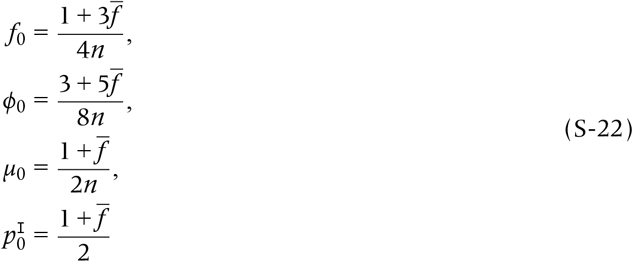

(because coalescence between offspring born in a patch aged *τ* = 0 may occur only if they share the same mother 1 / *n*). Also, multiplying *π_τ_* = *π_τ_*_−1_ ∙ (1 − *e*) with the recursions for (*f_τ_*, *ϕ_τ_*, *μ_τ_*, 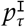) (Eqns (S-10) to (S-13)) and then summing up both sides over *τ* = 1, 2,…,supplies:

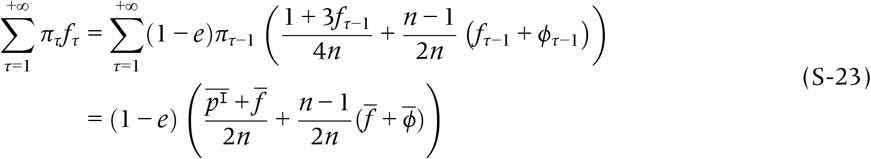

whereas the LHS is 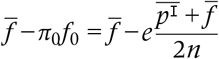. Using the similar algebra for (*ϕ*, *μ*, 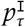), we get a closed relation for 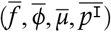:

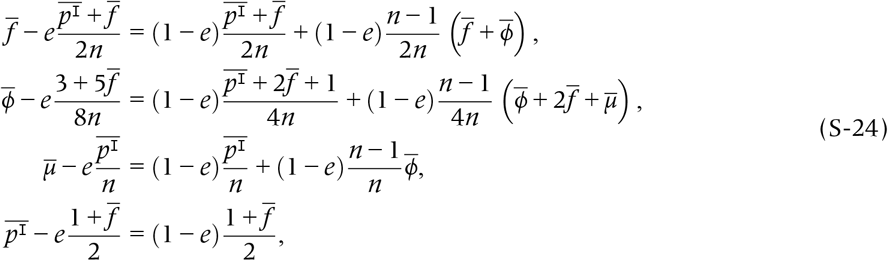

the last line of which implies 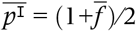, which further implies 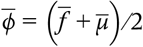. With some arrangement, we have:

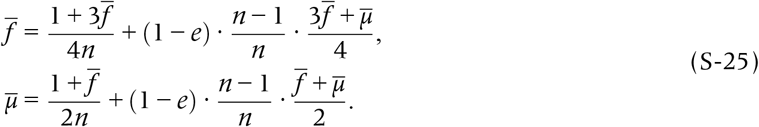

In a vector form,

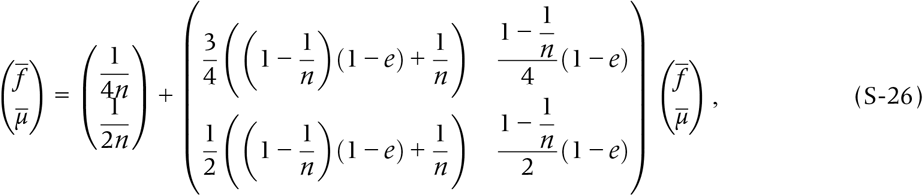

which gives:

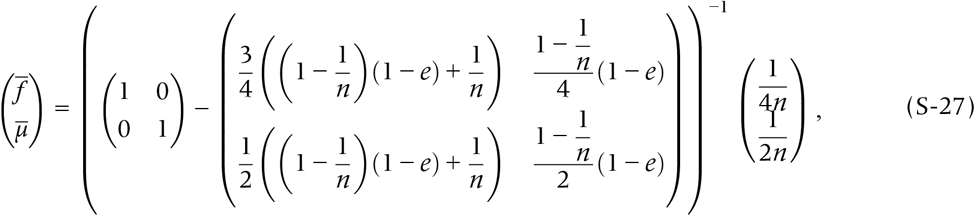

which is solved by:

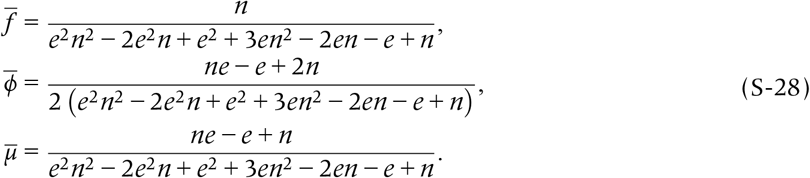

Substituting 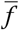 into the following equations gives the average values of *p*’s which read:

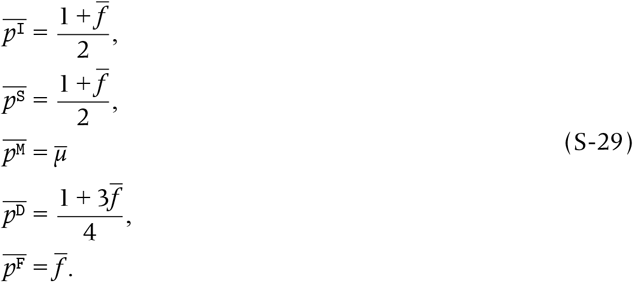

## 5 Fitness subcomponents

Let us focus on a patch aged *τ*. We denote the mutant sex allocation on the focal patch by *x_τ_*, the average sex allocation on the same patch by *y_τ_*, and the wild type sex allocation on the patch aged *τ* by *z_τ_*. In that patch, a focal female produces *J* (1 − *x_τ_*) of females (where *J* is the number of eggs per capita), who mate with the males born on the same patch. If that patch is persistent (with a probability of 1 − *e*), female offspring either disperse to recolonize empty patches with a probability of *d*, or else remain on their natal patch with a probability of 1 − *d*; if they do not disperse, they compete for reproduction on the patch against *J* (1 − *d*) (1 − *y_τ_*) of non-dispersing females (and therefore the factor *J* (1 − *d*) is cancelled out). If a patch is not persistent (with a probability of *e*), all females disperse for empty patches (*e* of the whole patches), and such dispersed offspring (*J* (1 − *x*_τ_)((1 − *e*) *d* + *e*) compete against on average 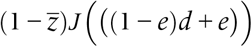 of female offspring (and therefore *J* (1 − *e*)*d* + *e* is cancelled out).

The daughter-fitness of a focal individual inhabiting on the focal patch 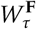 is therefore given by:

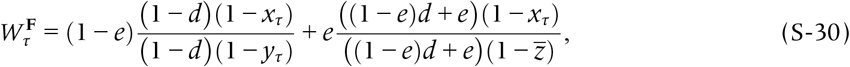

where we have written the spatial averaged sex allocation for 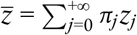, and the son-fitness of the focal adult female inhabiting in a patch aged *τ*, 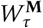, is given by:

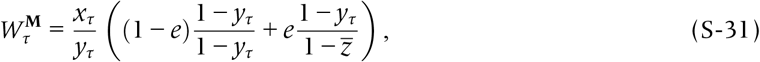

where we have eliminated the cancelling factors. Note that these fitness functions are defined such that neutrality (**x**= **y**= **z**) leads to 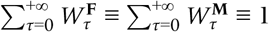.

Though we presumed in the main text that catastrophic extinction of patches occur after reproduction, but Eqns (S-30) and (S-31) can also be hold in other situations; for example: (i) a fraction 1 − *d* of females stays on their natal patch, and a fraction *d* disperses in the both persistent and non-persistent patches, (ii) all females stay on persistent patches, while all females disperse on non-persistent patches, and (iii) a fraction 1 − *d* of females stay and a fraction *d* disperse on persistent patches, while no females survive on non-persistent (extinct) patches. In all cases, the dispersal parameter *d* is cancelled out, and the fitness functions for daughters and sons are simplified to Eqns (S-30) and (S-31), respectively.

We can write the invasion fitness subcomponents in a general form as:

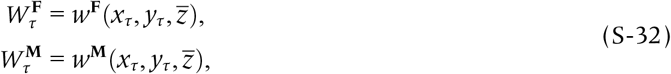

with

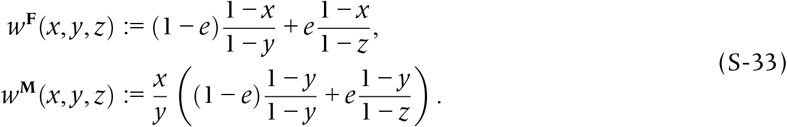

It is of use to write down the derivatives:

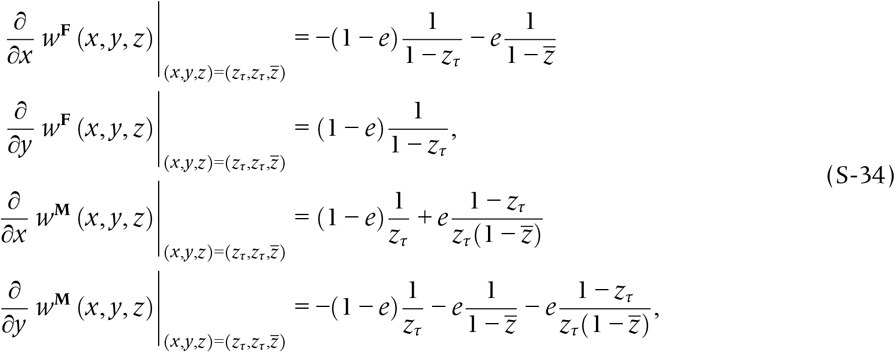

where note that the derivatives are evaluated at 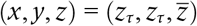. These quantities will be used below to assess the direction of selection under weak selection (Taylor & Frank 1996; Frank 1998; Rousset & Billiard 2000; Rousset 2004; Taylor et al. 2007).

## 6 Total fitness

Summing up the daughter- and son-mediated fitness functions (per capita) each multiplied by the class reproductive values, averaged over the patch-age distribution, obtains the total invasion fitness:

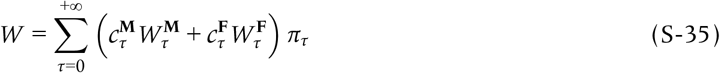

(Bulmer 1994; Taylor & Frank 1996; Frank 1998; Rousset 2004; Taylor et al. 2007; Lehmann & Rousset 2010). In particular, if the stratetegy is **z** = (*z*_E_, *z*_N_) (with a distribution *e*: 1 − *e*), *W* simplifies to:

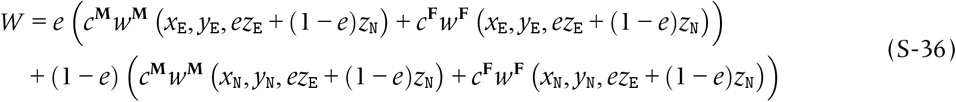

where the subscript _E_ accounts for disperser (Emigrant) females (hence inhabiting on the patch aged *τ* = 0), while _N_ accounts for non-disperser females (hence on the patch aged *τ* > 0). Also we have here made it explicit that 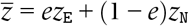. Also, class reproductive values (Taylor 1990; Caswell 2001) are denoted *c*^**M**^ for male and *c*^**F**^ for females.

## 7 Reproductive value

As the patch-age generates no difference in reproductive capacity, the class reproductive values are independent of patch ages and are fully determined by the ploidy: *c*^**M**^ = *c*^**F**^ = 1 /2 for diploidy, and *c*^**M**^ = 1 /3, *c*^**F**^ = 2 /3 for haplodiploidy (Taylor 1990; Caswell 2001).

## 8 Selection gradient

We outline the analyses for the general case in which the trait to evolve is patch age-specific sex ratio sorted as **z**= (*z*_0_, *z*_1_,…, *z_τ_*,…), where *z_τ_* represents the sex ratio strategy of a female breeding on a patch aged *τ*. The selection gradients for dispersers’ and non-dispersers’ strategy *z*_E_ (E for emigrants) and *z*_N_ (N for non-disperser) are, using the neighbor-modulated fitness approach (Taylor & Frank 1996; Frank 1998; Rousset & Billiard 2000; Rousset 2004; Taylor et al. 2007), given by:

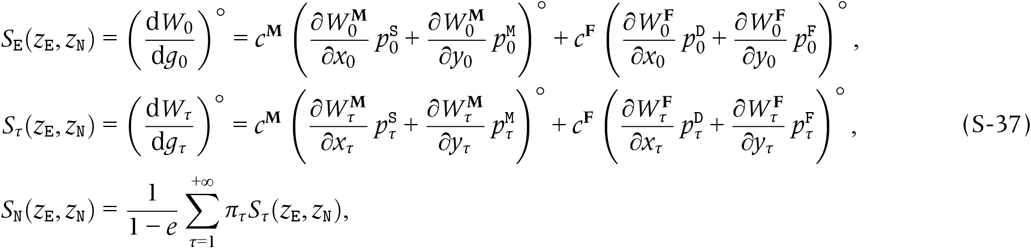

where ° represents neutrality, (i.e., the derivatives are evaluated at **x**= **y**= **z**). *g_τ_* represents the genic value of a gene sampled from a locus (denoted 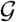, that encodes the sex allocation) of a female offspring (Falconer 1975; Grafen 1985; Bulmer 1994; Taylor & Frank 1996; Frank 1998; Gardner et al. 2009). Also, *p*-values are the consanguinities of an adult female with an corresponding offspring sharing the same patch (age *τ*): S designates her own son, M male offspring, D her own daughter, and F female offspring, respectively.

## 9 Unconditional strategy

When females exhibit unconditional strategy (i.e., *z_τ_* ≡ *z*_U_ for all *τ* ≥ 0), the selection gradient reads:

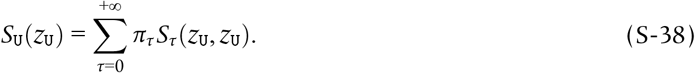

ESS allocation simplifies down to:

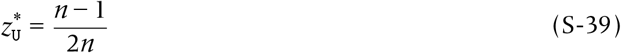

for diploids, and:

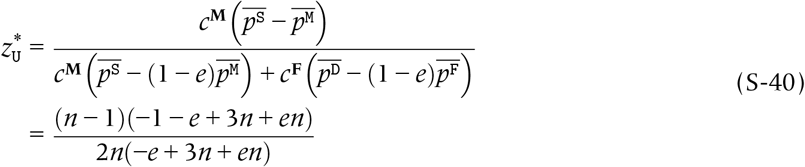

for haplodiploids. We therefore recover Gardner *et al.*’s (2009) results by replacing *e* with 1 − (1 − *d*)^2^ (where *d* is female-dispersal rate after mating).

## 10 Dispersers’ strategy

Higher male allocation is favored for a disperser female if:

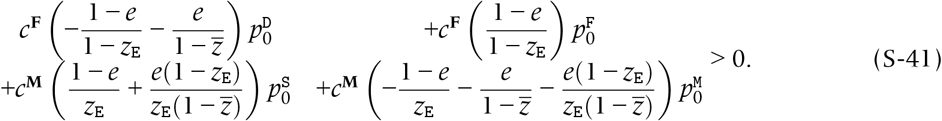

If we divide both sides by 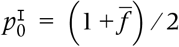 (the consanguinity of a mother to herself), we get the Hamilton’s rule of the main text, after clearing the fractions.

## 11 Non-dispersers’ strategy

Higher male allocation is favored for a non-disperser female if:

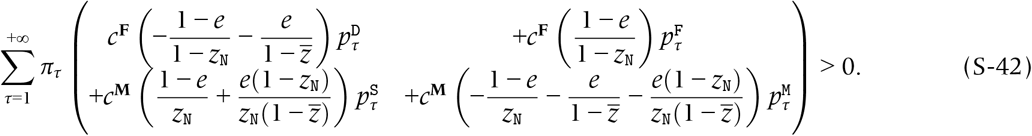

The quantities inside the bracket are dependent on patch age *τ* only through consanguinity, *p*-values. Therefore, what matters is the average values of *p*’s (minus *ep*_0_), given that she is in a patch aged *τ* ≥ 1:

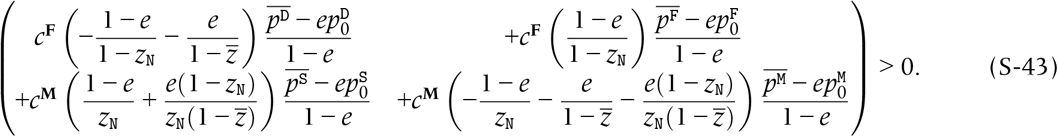

Dividing both sides by 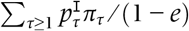 (average consanguinity of a mother to herself in a patch aged *τ*), we get Hamilton’s rule of the main text,after clearing the fractions.

## 12 Evolutionary outcomes

We obtained evolutionary outcomes 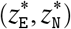 by nullifying the Hamilton’s rules, and we call the pair of the evolutionary outcomes as cESSs (the candidate ESSs; Maynard Smith & Price 1973; Hofbauer & Sigmund 1990; Geritz et al. 1998).

### Sensitivity to the number of females ovipositing on a patch, *n*

The cESSs monotonically increase with *n*, and eventually lead to Fisherian sex ratio 1/2 with *n* → +∞ (Supplementary Fig 3-1A). When we compare 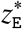 and 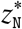, we found that 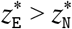 always holds true for diploids. The cESSs generally increase with decreasing patch extinction rate *e* (Supplementary Fig 3 - 1A; but except large *n* for haplodiploidy), which is likely to be because local competition between related females increases with smaller *e* (Bulmer 1986; Taylor 1988b; Frank 1998; Gardner et al. 2009).

However, we found the predicted patterns complicated for haplodiploids. For intermediate or high extinction rates (for example, *e* = 0.5 or 0.8), 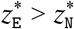 is also favoured. In contrast, for small *e*(= 0.2), small *n* favours 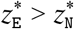 while the opposite 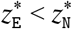 appears to occur when *n* is larger. This trend can be explained by relatedness asymmetry for haplodiploid sex determination. For dispersers, inbreeding increases relatedness of mothers to their daughters, but does not relatedness to their sons, which favours a more female-biased sex ratio for haplodiploid than diploid species (Supplementary Fig 3 - 1A; Frank 1985; Herre 1985). This trend is remarkable with smaller *e* (i.e., higher inbreeding rates). For non-dispersers, mothers are related not only with their own offspring but also with offspring produced by other mothers on the same patch. This leads to almost identical cESSs between diploids and haplodiploids (Supplementary Fig 3 - 1A; Hamilton’s rules for non-dispersers are exactly identical for diploids and haplodiploids). Consequently, 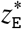 is predicted to be more female biased than 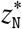, when *e* is small and *n* is large.

Overall, the cESSs are predicted to increase with *n*, and decrease with inbreeding rates, in agreement with to the prediction by a theoretical model assuming that females are able to adjust their offspring sex ratio according to the number and kinship of females laying eggs on a patch (Gardner & Hardy 2020).

### Sensitivity to relatedness between offspring on a patch

In the present model, the influences of the number of mothers and relatedness between mothers on a patch can be summarized to one parameter, relatedness between offspring on a patch. We define relatedness between female offspring on a patch produced by dispersers and non-dispersers as:

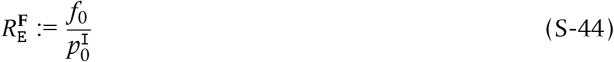

and

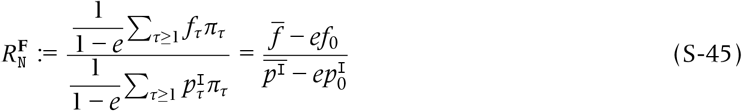

respectively. We first assessed the effects of *n* (the number of mothers ovipositing on a patch) on the relatedness coefficients, and found that relatedness coefficients decrease with *n* and that 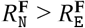 (Supplementary Fig 3 - 2), which is because non-disperser females are more likely to be related with neighboring females than are disperser females (El Mouden & Gardner 2008; Wild & Fernandes 2009).

By plotting the cESSs against the relatedness (by tuning *n*), we found that more female-biased sex ratios are favoured with increasing relatedness between offspring (Supplementary Fig 3 - 1B). This negative relationship is predicted for both dispersers and non-dispersers, although the detailed patterns depend on the difference of ploidy. While 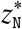 is similar for diploids and haplodiploids, 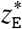 is more female biased for haplodiploid than diploids especially with smaller 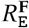 and smaller *e*. Consequently, cESSs for dispersers and non-dispersers are predicted to switch at an intermediate value of relatedness for haplodiploid species (Supplementary Fig 3 - 1B).

By plotting the cESSs in terms of relatedness, we can separately investigate the effects of relatedness and local competition between relatives (Cooper et al. 2018). Here, the scale of competition equals 1 − *e*, which is the probability that two randomly chosen females laying eggs on a patch are derived from the same patch (Frank 1998). We found that less female-biased sex ratios are predicted with higher local competition (smaller *e*) for both dispersers and non-dispersers (Supplementary Fig 3 - 1B; see also Gardner et al. 2009, in which the scale of competition is (1 − *d*)^2^, where *d* is female dispersal rate). In the natural populations, the effects of relatedness and local competition between relatives are likely to influence the evolution of sex ratio, with its extent dependent upon life history details, such as population structure and whether females can assess if they are with closer relatives (Frank 1985; Frank 1986; Frank 1998; Bulmer 1986; Taylor 1988b; Gardner et al. 2009; Lehmann & Rousset 2010; Cooper et al. 2018; Gardner & Hardy 2020).

**Supplementary Fig 3-1:**
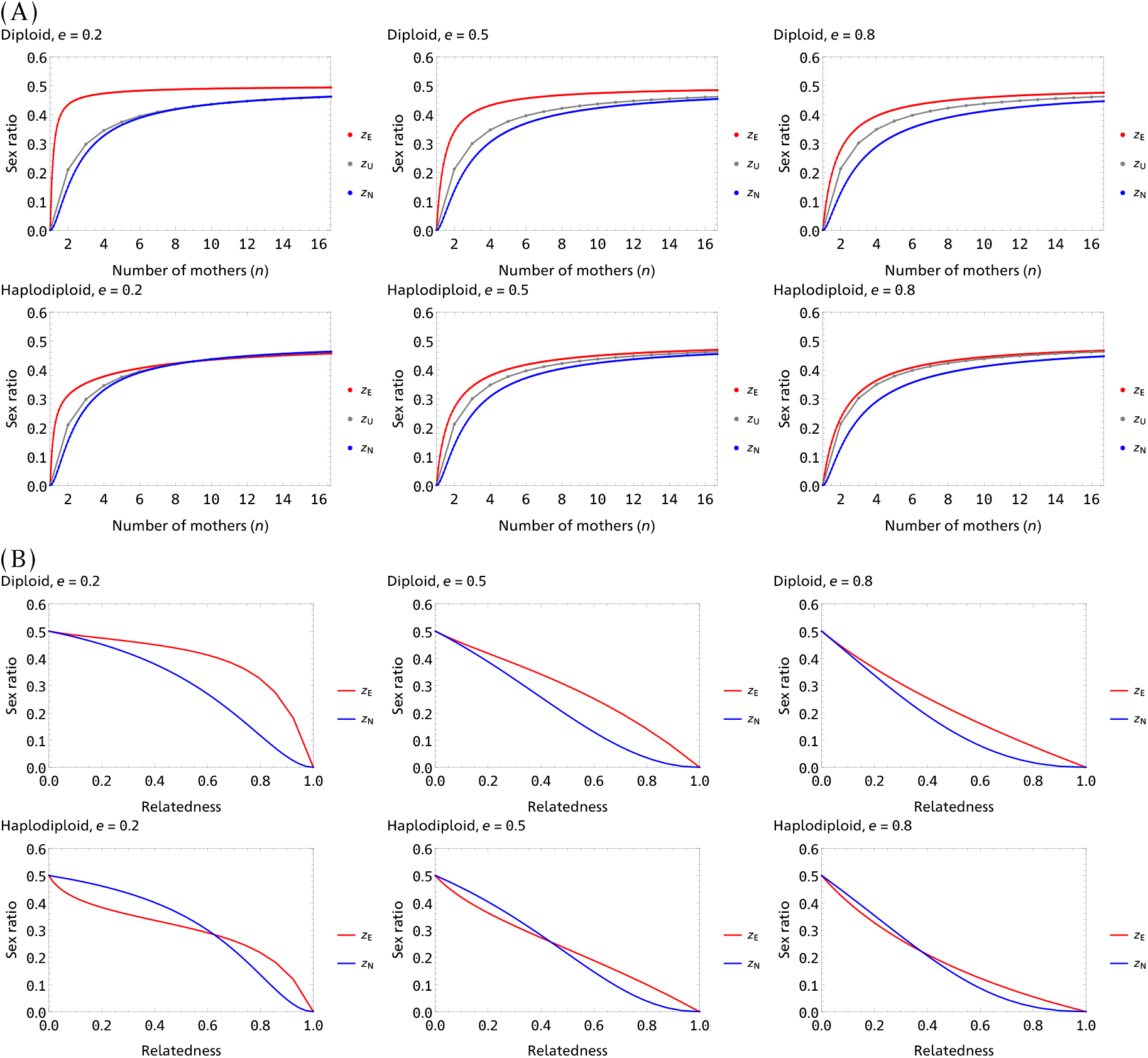
Predicted sex ratio (proportion sons) plotted against the number of females ovipositing on a patch (2; panel A) and against the relatedness coefficient for female offspring on a patch 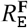 for dispersers and 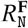 for non-dispersers (panel B).

**Supplementary Fig 3-2:**
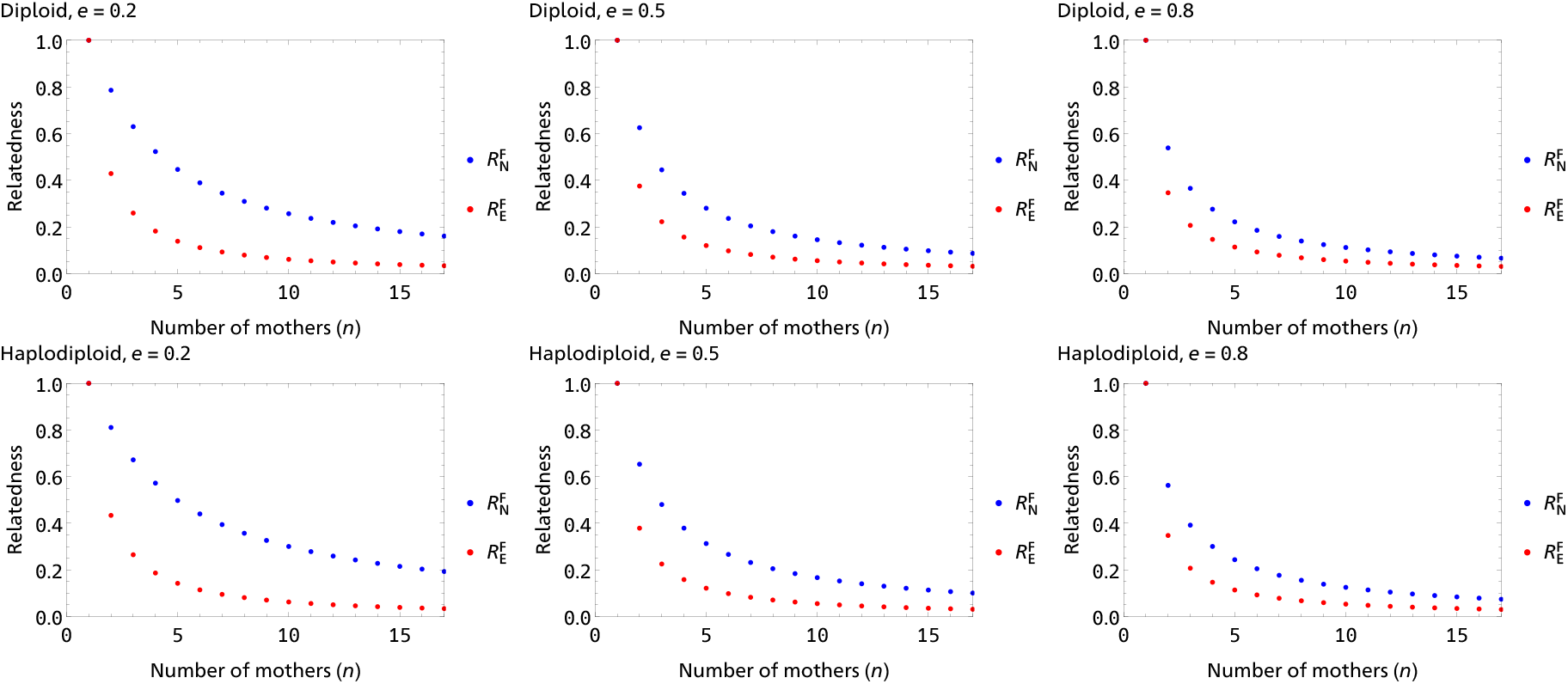
Relatedness coefficients plotted against *n*.

